# Xylulose 5-Phosphate/Phosphate Translocator Mediates Carbon Allocation for Anabolic Metabolism in the Chlamydomonas Chloroplast

**DOI:** 10.64898/2026.02.27.708587

**Authors:** Anbang Wu, Nicole Linka, Yue Wu, Luca Frederik Schneider, Sijie Xu, Qingling Kong, Yonghong Bi, Arthur R. Grossman, Weichao Huang

## Abstract

Metabolite movement into chloroplasts is essential for sustaining chloroplast anabolic metabolism and cellular growth, and yet how the specific factors/transporters that enable import and support metabolism under heterotrophic conditions (in the dark in the presence of fixed carbon) remain poorly understood. Here, we identify CreTPT10, a chloroplast envelope-localized transporter in the unicellular green alga *Chlamydomonas reinhardtii* (Chlamydomonas throughout) that, of the substrates tested, has the highest specificity for xylulose 5-phosphate (X5P). We also demonstrated that the loss of CreTPT10 caused a pronounced reduction in growth and respiratory activity in the dark, whereas no growth defects were observed in the *tpt10* mutants when they were maintained in the light or under nutrient-limiting conditions. Furthermore, dark-grown *tpt10* mutants exhibited markedly reduced levels of lipids, nucleotides, isoprenoids, and aromatic secondary metabolites, accompanied by coordinated repression of genes encoding enzymes associated with chloroplast-localized biosynthetic pathways. This metabolic suppression extended beyond the chloroplast, as genes associated with the mitochondrial respiratory chain and cell cycle progression were markedly downregulated in darkness. Together, these findings indicate that X5P import via CreTPT10 is critical for sustaining chloroplast anabolic metabolism and functionally coordinates chloroplast and mitochondrial energy metabolism to support heterotrophic growth.

## Introduction

Microalgae represent a highly promising source of high-value bioactive compounds, with broad applications in human nutrition, food, cosmetics, pharmaceuticals, bioplastics, and energy (Borowitzka, 2013; Levasseur et al., 2020). Notably, many of these compounds or their biosynthetic precursors, including photosynthetic pigments, amino acids, lipids, isoprenoids/terpenoids, etc., are synthesized in the chloroplast (Ohlrogge and Browse, 1995; Neuhaus and Emes, 2000; Tanaka and Tanaka, 2007; Maeda and Dudareva, 2012; Ruiz-Sola and Rodríguez-Concepción, 2012). Maintaining an integrated metabolic network in the cell requires extensive metabolite exchange between the chloroplast and cytosol, driven by specific organic carbon transport systems. Moreover, microalgae often exhibit diverse modes of nutrition, such as photoautotrophy, heterotrophy, and mixotrophy, implying that the metabolite transport systems on the chloroplast envelope are subject to active regulation and substantial remodeling in response to shifts in trophic status. Chloroplasts in some microalgal species, such as *Chlamydomonas reinhardtii* (hereafter Chlamydomonas), retain a fully assembled and functional photosynthetic electron transport chain even in darkness. Photoautotrophy (light and no fixed carbon) relies heavily on the export of photoassimilates (e.g., carbon compounds and reducing equivalents) from chloroplasts to the cytosol (and then to other subcellular compartments), whereas heterotrophic growth depends on the import of organic carbon into chloroplasts to fuel chloroplast-localized biosynthetic pathways. Despite the central roles of these transport systems in sustaining metabolic flux and regulation, those transporters in algae responsible for trafficking specific metabolites across the chloroplast envelope remain largely unidentified. This knowledge gap significantly hinders the application of synthetic biology approaches to enhance production of valuable compounds in these organisms.

Plastid phosphate translocators (pPTs) are the major transporters on the inner envelope membrane of the chloroplast. They function as antiporters that exchange inorganic phosphate (Pi) for phosphorylated C3, C5, or C6 sugars (Weber et al., 2005). Based on substrate specificity, pPTs are classified into four functional classes: triose phosphate/phosphate translocator (TPT), glucose 6-phosphate/phosphate translocator (GPT), phosphoenolpyruvate/phosphate translocator (PPT), and xylulose 5-phosphate (X5P)/phosphate translocator (XPT) (Flügge et al., 2003). Among these transporters, certain members play a crucial role in the export of photosynthetically derived carbohydrates from the chloroplast to the cytosol. In Chlamydomonas, the TPT homolog CreTPT3 is indispensable and serves as the primary conduit that brings both carbon and reductant across the chloroplast envelope (Huang et al., 2023). In contrast, in vascular plants, the loss of TPT function can be compensated through the export of maltose derived from starch degradation via the maltose exporter (MEX) (Niittylä et al., 2004; Cho et al., 2011). In addition, in *Arabidopsis* (*Arabidopsis thaliana*), XPT can also export fixed carbon from the chloroplast, thereby partially compensating for the loss of TPT (Hilgers et al., 2018). Furthermore, in C_4_ and CAM plants, PPTs export phosphoenolpyruvate from the chloroplast to the cytosol, where it serves as a substrate for CO_2_ fixation (Fischer et al., 1997; Häusler et al., 2000; Majeran et al., 2008).

Besides these export pathways, chloroplasts depend on the import of cytosolic metabolites to sustain their anabolic metabolism, especially when the cells are maintained in the dark. In a non-photosynthetic diatom, four plastid-localized TPT homologues have been identified, three of which have been demonstrated in vitro to mediate the transport of phosphorylated metabolites—including triose phosphates, phosphoenolpyruvate, and 3-phosphoglycerate. However, their physiological roles in vivo remain unresolved, although it has been hypothesized that these putative TPTs facilitate the import of cytosol-derived carbon and energy sources into the plastid to support the metabolic activities retained in the non-photosynthetic organelle (Kamikawa et al., 2017; Moog et al., 2020). With the import of cytosolic glucose 6-phosphate into the stroma of non-photosynthetic plant plastids, GPTs support starch biosynthesis, the oxidative pentose phosphate pathway, and fatty acid synthesis (Kammerer et al., 1998; Niewiadomski et al., 2005; Rolletschek et al., 2007; Zheng et al., 2018). In C_3_ plants, the PPTs supply plastids with cytosolic phosphoenolpyruvate, thereby fueling aromatic amino acid biosynthesis in leaves as well as glycolytic metabolism and lipid accumulation in seeds (Fischer et al., 1997; Streatfield et al., 1999; Staehr et al., 2014; Tang et al., 2022). By contrast, the physiological role of XPT remains less clear, as single *Arabidopsis* mutants exhibit no obvious phenotype (Hilgers et al., 2018), although XPT is thought to import cytosolic X5P into plastids, supporting the pentose phosphate pathway under conditions of high metabolic demand (Eicks et al., 2002; Weber et al., 2004).

Collectively, pPTs are recognized as an integrated transport system that supports plastid anabolic metabolism and redox homeostasis, although this framework has been established largely based on studies of vascular plants. The mechanisms by which microalgal chloroplasts import and utilize cytosolic carbon are not as well understood. Given that many microalgae are capable of heterotrophic or mixotrophic (light plus organic carbon source) growth, elucidating plastid carbon import pathways in these organisms represents a critical gap in our current understanding of plastid metabolism.

Here, we used Chlamydomonas, a widely adopted model alga to investigate chloroplast organic carbon import under heterotrophic conditions (dark plus acetate). We showed that *CreTPT10* was specifically induced under dark conditions and that the CreTPT10-VENUS protein fusion was localized to the chloroplast envelope. Loss of CreTPT10 caused growth defect and metabolic changes under heterotrophic conditions, but not under photoautotrophic conditions; these differences were accompanied by reduced respiration. We also reported the impact of this transport defect on both the transcriptome and metabolome of the cells. Disruption of this transporter triggered pronounced transcriptomic and metabolomic changes, particularly affecting chloroplast anabolic and respiratory pathways. Using a combination of genetic, biochemical and multi-omics approaches, we demonstrated that CreTPT10 functions as a chloroplast envelope importer, preferentially transporting X5P to sustain chloroplast anabolic metabolism and mitochondrial respiration when photosynthetic carbon supply is absent.

## Results

### Expression Patterns and Subcellular Localization of CreTPT10

To identify translocators potentially crucial for sustaining chloroplast activity under heterotrophic conditions in Chlamydomonas, we analyzed the diurnal expression profiles of four *pPT* genes (*CreTPT2*, *CreTPT3*, *CreTPT10*, and *CreCGL51*) using published datasets from Dupuis et al. (2025). As shown in Figure 1A, among the four pPT-encoding genes, *CreTPT10* exhibited the most pronounced responses to dark-light transitions. *CreTPT10* transcript levels were rapidly downregulated following the transition from darkness to light, declining to near the detection limit, suggesting that light may act as a repressive signal for its expression (Figure 1B). In contrast, following the light-to-dark shifts, *CreTPT10* was markedly induced by darkness, exhibiting the highest fold increase in abundance (4–11 fold) (Figure 1B). Based on the phylogenetic analyses of plant and algal pPTs, CreTPT10 grouped within the putative PPT clade together with CreTPT2 (Supplemental Figure 1). Moreover, we confirmed that CreTPT10 fused to VENUS localized to the chloroplast envelope (Figure 1C). This distinct expression pattern and its location in the cell suggest CreTPT10 may fulfill a specific role during the dark phase, potentially providing metabolites for chloroplast function under heterotrophic conditions.

**Figure 1.**
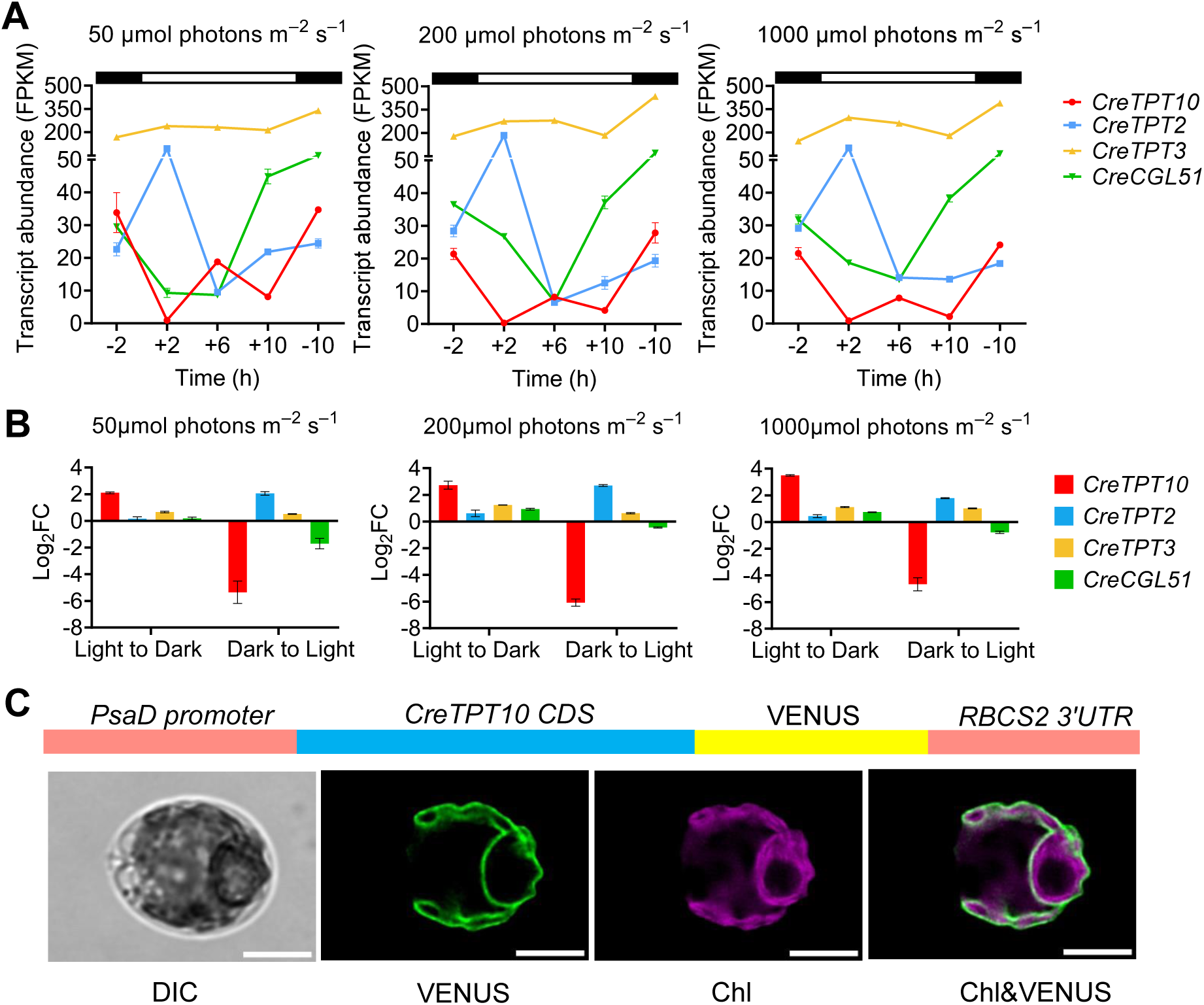
Expression of *pPT* genes and subcellular localization of CreTPT10 in Chlamydomonas. **(A)** Expression patterns of *CreTPT10*, *CreTPT2*, *CreTPT3*, and *CreCGL51* during a 12 h light/12 h dark diurnal cycle under light of 50, 200, and 1000 μmol photons m^-2^ s^-1^. Black and white bars above the plots indicate dark and light phases, respectively. Transcript levels are shown as FPKM values. Transcriptomic data were extracted from Dupuis et al. (2025). **(B)** Transcriptional responses of *CreTPT10*, *CreTPT2*, *CreTPT3*, and *CreCGL51* to light-to-dark transitions at light intensities of 50, 200, and 1000 μmol photons m^-2^ s^-1^. For the dark to light transition, log_2_FC was calculated by normalizing transcript abundances at +2 h (2 h in light) to the mean abundance at -2h (10 h in dark); for the light to dark transition, log_2_FC was calculated by normalizing transcript abundance at -10 h (2 h in dark) to the mean abundance at +10 h (10 h in light). Data were extracted from Dupuis et al. (2025). **(C)** Cellular localization of CreTPT10. A schematic of the construct encoding the CreTPT10-VENUS fusion protein is shown above the cell images and described in Methods. Representative differential interference contrast (DIC), VENUS fluorescence (VENUS), chlorophyll autofluorescence (Chl), and merged (Chl & VENUS) images are shown. Scale bars, 5 μm.

### Generation of *tpt10* Knockout Mutants and Their Effects on Cell Growth and Respiration

To explore the role of CreTPT10 in trafficking carbon between the chloroplast and cytosol in darkness, we generated knockout mutants of *CreTPT10* by CRISPR/Cas9-mediated gene editing. CRISPR/Cas9 editing was used to disrupt the *CreTPT10* locus while simultaneously integrating a hygromycin-resistance marker gene (*AphVII*) at the edited site (Figure 2A). We obtained 3 independent, putative transformants knockouts for *CreTPT10* (*tpt10a*, *b*, and *c*), with the marker gene inserted into exon 9 (Supplemental Figure 2). The *CreTPT10* transcript levels were strongly reduced in all three edited *tpt10* mutants compared with the level in WT cells (Figure 2B), confirming successful gene disruption.

**Figure 2.**
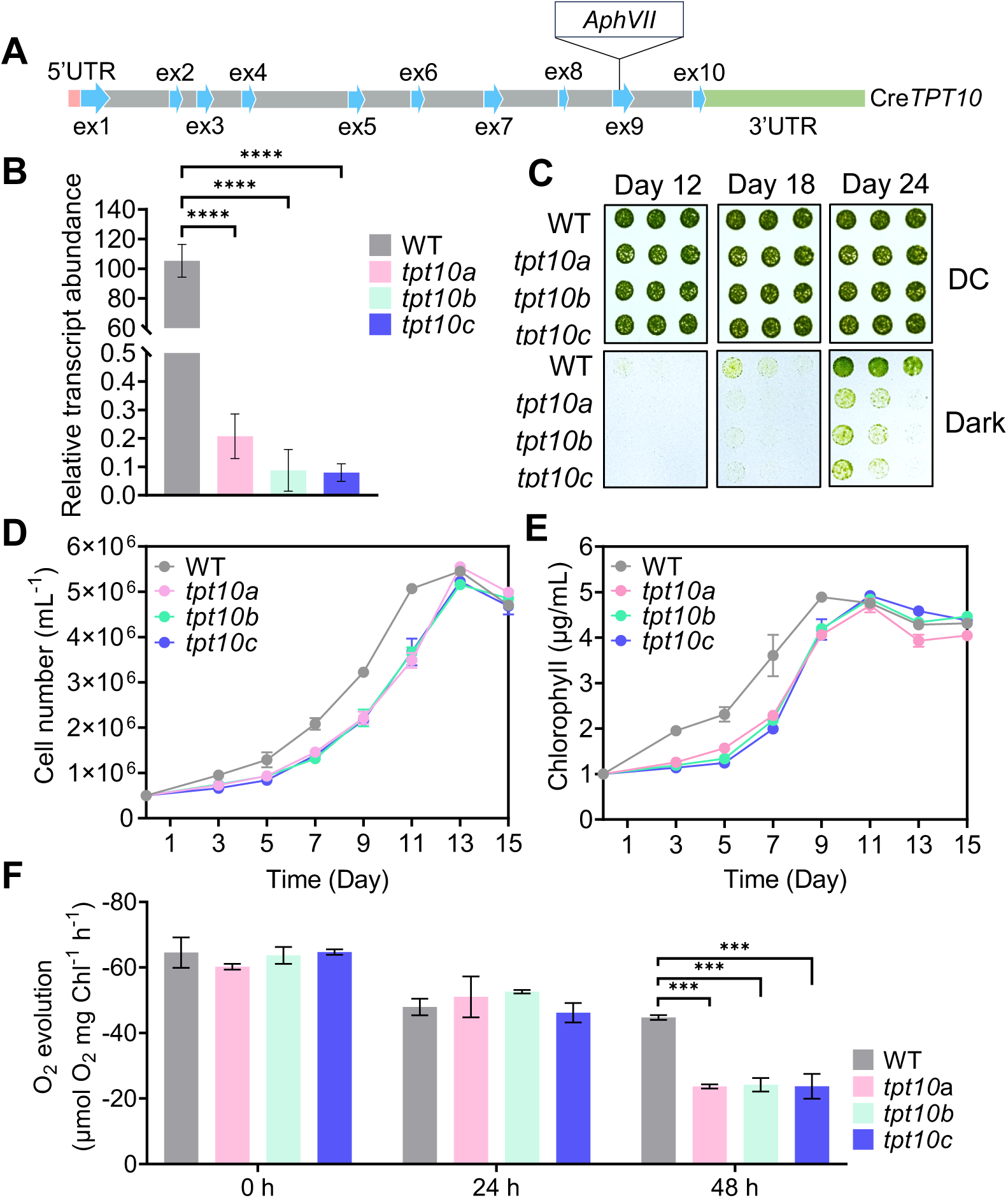
Characterization of *tpt10* knockout mutants in Chlamydomonas. **(A)** Schematic representation of *CreTPT10* and Cas9 target sites, with the edited site located in exon 9. **(B)** *CreTPT10* transcript abundance in WT and three independent knockout strains (*tpt10a*, *tpt10b*, *tpt10c*) measured by RT-qPCR. The asterisks represent statistically significant differences determined by one-way ANOVA: ****P < 0.0001. **(C)** Growth analysis of WT and *tpt10* mutants (*tpt10a*, *tpt10b*, *tpt10c*) on TAP agar plates. Cells were spotted on agar plates containing TAP medium and maintained under diurnal cycle (DC, 30 μmol photons m^-2^ s^-1^) or extended darkness for 12, 18, and 24 d. The dilution series used was 4, 2, and 1 μg mL^-1^ chlorophyll equivalent (left to right). **(D and E)** Cell growth based on cell number **(D)** or chlorophyll content **(E)** at various times after the cells were shifted from LL to darkness for up to 15 d. Cultures were initially grown in TAP medium at 5 × 10^5^ cells mL^-1^ or 1 μg chlorophyll mL^-1^ under LL before transfer to the dark. Each curve represents the arithmetic mean (± SEM) of three independent experiments. **(F)** Respiration during darkness. Respiration rates of WT and three independent *tpt10* mutants were measured at 0 h (prior to transfer to darkness) and after 24 and 48 h in the dark. Data represent the arithmetic mean (± SEM) of three independent experiments (n = 3). Asterisks indicate statistically significant differences determined by one-way ANOVA: ***P < 0.001.

To elucidate the physiological roles of CreTPT10, we examined the growth of WT and *tpt10* mutants on solid agar plates under various conditions, including darkness, continuous low light (LL), continuous high light (HL), diurnal cycle (DC), and nitrogen-deficient conditions (−N) (Figure 2C; Supplemental Figure 3). Notably, a severe growth defect was observed in all three *tpt10* mutants (*tpt10a*, *b*, and *c*) under dark conditions, whereas no significant differences in the growth of the mutants relative to WT cells were observed under other tested conditions (Figure 2C; Supplemental Figure 3). Consistent with their growth phenotype on solid medium, all three *tpt10* mutants also exhibited significant growth impairment in liquid medium compared to the WT cells (Figures 2D and 2E), although all strains ultimately reached similar cell densities at stationary phase. Additionally, all three *tpt10* strains exhibited markedly reduced respiration rates compared with WT cells after 48 h in darkness (Figure 2F). Taken together, these results indicate that the activity of the CreTPT10 transporter in Chlamydomonas is specifically required to sustain growth and respiration under prolonged darkness.

### Biochemical Characterization of CreTPT10

The transport activity of CreTPT10 was characterized using proteoliposome-based uptake assays after reconstitution with recombinant protein. To this end, the coding sequence of the mature CreTPT10 protein was codon-optimized for heterologous expression in yeast and the N-terminal transit peptide was replaced by a His tag. Successful expression and membrane localization of His-CreTPT10 were confirmed in enriched membrane fractions by immunodetection using an anti-His antibody (Figure 3A). Functional analyses demonstrated that His-CreTPT10 reconstituted into liposomes functions as an antiporter. Time-course experiments revealed a time-dependent uptake of inorganic phosphate (Pi) when Pi was present as a counter-substrate inside the lipid vesicles. In contrast, no measurable Pi uptake was detected in the absence of an internal exchange substrate, confirming the strict counter-exchange mechanism of CreTPT10 (Figure 3B). Kinetic characterization based on Michaelis–Menten analysis yielded an apparent K_M_ value for Pi of 2 mM (Figure 3C). This value is approximately twofold higher than that reported for other members of the Chlamydomonas pPT family. Notably, the determined K_M_ exceeds the estimated physiological stromal Pi concentration, suggesting that Pi may not represent the preferred substrate of His-CreTPT10. To further define substrate specificity, Pi uptake was quantified in proteoliposomes preloaded with various potential counter-substrates (Figure 3D). The highest Pi uptake rates were observed when liposomes were internally loaded with X5P (150%) or triose phosphates (110%), relative to Pi/Pi homo-exchange. Additional substrates supported Pi exchange to a lesser extent. These findings indicate a broader substrate spectrum with a clear preference for phosphorylated pentose- and triose-phosphate intermediates. To obtain kinetic parameters for X5P and triose phosphates, competitive inhibition assays were performed by measuring Pi/Pi exchange rates in the presence of increasing external concentrations of these metabolites. Both X5P and triose phosphates inhibited Pi/Pi exchange in a competitive manner (Figure 3E). The inhibition constants (K_i_) were determined to be 0.5 µM for X5P and 0.85 µM for triose phosphates. These K_i_ values are approximately 4,000-fold lower than the K_M_ for Pi, indicating substantially higher affinities for X5P and triose phosphates. These in vitro data strongly suggest that CreTPT10 functions predominantly in the exchange of X5P and triose phosphates across the chloroplast inner envelope membrane under physiological conditions.

**Figure 3.**
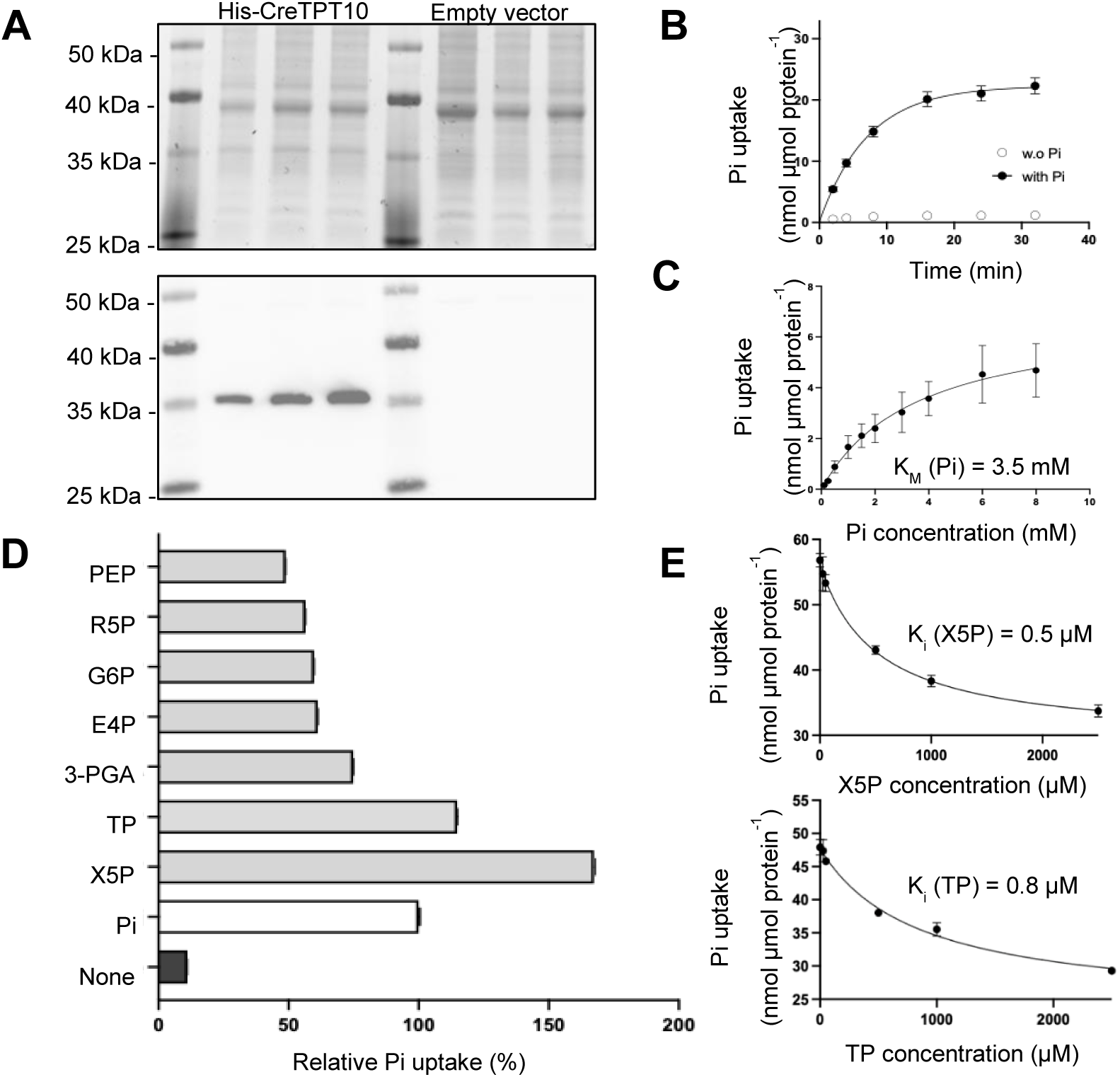
Heterologous expression of CreTPT10 in yeast and in vitro measurements of transport activities. **(A)** Production of His-CreTPT10 mature protein in yeast. Total membrane fractions from cells expressing His-CreTPT10 or the empty vector were analyzed by protein-stained SDS-PAGE (top) and anti-His immunoblotting (bottom). The calculated molecular mass of mature His-CreTPT10 is 39.3 kDa. **(B)** Time course of [^32^P]-Pi uptake (0.25 mM external concentration) into liposomes reconstituted with His-CreTPT10 and either lacking internal substrate (○) or preloaded internally with 25 mM Pi (●). **(C)** Michaelis–Menten constant (K_M_) of His-CreTPT10 for Pi determined from Pi/Pi exchange rates measured at increasing external Pi concentrations (0.25–8 mM), while maintaining an internal Pi concentration of 25 mM. The K_M_ value was obtained by non-linear regression analysis. **(D)** Substrate specificity was determined by measuring the [^32^P]-Pi uptake (0.25 mM external concentration) into liposomes reconstituted with recombinant His-CreTPT10 and preloaded with various substrates (25 mM) or in the absence of any substrate. The velocity of Pi uptake after 2 min was determined. Relative uptake activities were compared with the Pi/Pi homo-exchange experiment, which was set to 100%. **(E)** The inhibition constant (Ki) of His-CreTPT10 for X5P (top) or triose phosphate (bottom) was determined from [^32^P]-Pi uptake at an external Pi concentration of 0.25 mM in exchange with 25 mM internal Pi, while increasing concentrations of X5P or triose phosphate (0.025–2.5 mM) were applied. The Ki values were calculated by non-linear regression analysis. Data in **(B–E)** are arithmetic means ± SEM from three independent experiments. Abbreviations: PEP, phosphoenolpyruvate; R5P, ribose 5-phosphate; G6P, glucose 6-phosphate; E4P, erythrose 4-phosphate; 3-PGA, 3-phosphoglycerate; TP, triose phosphate; X5P, xylulose 5-phosphate; Pi, inorganic phosphate.

### Metabolic Changes in *tpt10* Mutants after shifting the cells to the Darkness

To explore the metabolic impact of a CreTPT10 deficiency under light and dark conditions, we performed comprehensive metabolite profiling of WT cells and two independent *tpt10* mutants (*tpt10a* and *tpt10b*) during the transition from light (0 h) to darkness (4 h and 48 h). Quality control (QC) samples clustered tightly by Principal Component Analysis (PCA), indicating high analytical reproducibility across the acquisition batch (Supplemental Figure 4A). Based on positive- and negative-ion electrospray ionization (ESI) modes, 4,198 and 2,089 metabolic features were detected, respectively (Supplemental Data Set 1). Under light (0 h) and after 4 h of extended darkness, PCA showed no clear metabolic separation between WT cells and the *tpt10* mutants (ANOSIM, P > 0.05; Supplemental Figures 4B, 4C and 5; Supplemental Data Set 2). In contrast, after 48 h of extended darkness, PCA revealed a clear and statistically significant divergence of the *tpt10* metabolic profiles from those of WT cells (ANOSIM, R = 0.3703, P = 0.014; Supplemental Figures 4D and 5). Consistent with this separation, the number of differentially abundant metabolites (DAMs) increased markedly after 48 h in the dark, which included 70 metabolites with higher abundance and 134 with lower abundance in the mutants relative to WT cells (Supplemental Data Set 2).

After 48 h of extended darkness, 20 DAMs associated with fatty acid and lipid metabolites were identified, all of which were reduced in the *tpt10* mutants relative to WT cells (Supplemental Table 1). Notably, malonyl-CoA, which functions in the initial step of de novo fatty acid synthesis in chloroplasts, declined to 60–64% of the WT cell levels, accompanied by a reduction in the downstream β-ketoacyl intermediate 3-oxooctanoyl-CoA (Figures 4A-1 and 4B; Supplemental Figure 6; Supplemental Table 1). Consistently, the levels of downstream lipid products also declined, including 3 fatty acids (eicosapentaenoic acid, myristic acid, and γ-linolenic acid), 12 triacylglycerol species and three phosphatidylethanolamine species (Figures 4A-1 and 4B; Supplemental Figure 6; Supplemental Table 1). These coordinated decreases across fatty acid precursors, biosynthetic intermediates, and complex lipid species indicate a broad suppression of chloroplast fatty acid and lipid biosynthesis in the *tpt10* mutants during prolonged darkness.

**Figure 4.**
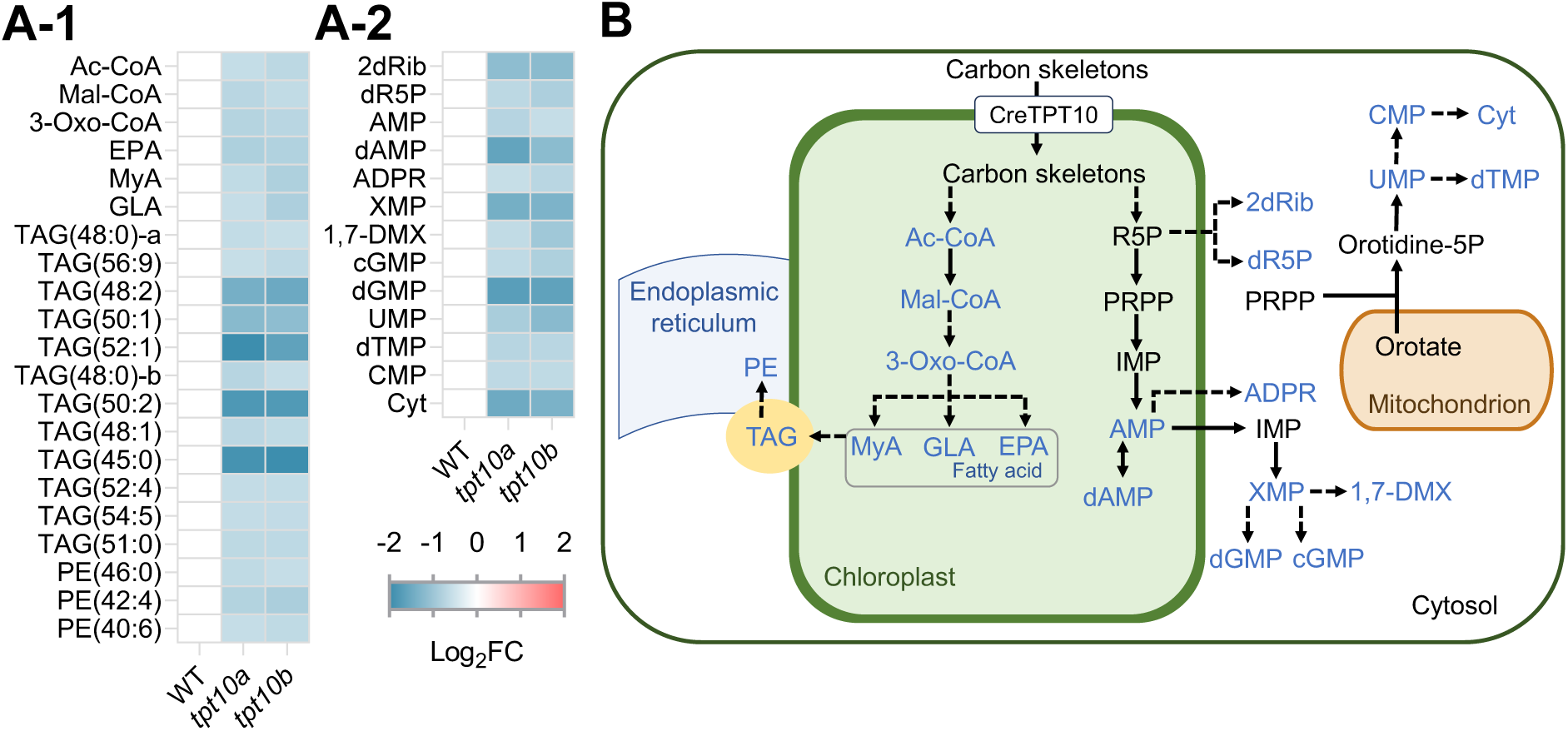
Changes in lipid/fatty acid- and nucleotide-related metabolites in WT, *tpt10a*, and *tpt10b* under extended darkness. **(A)** Heatmaps of significantly altered lipid/fatty acid- and nucleotide-related metabolites after 48 h of extended darkness, shown as log_2_-transformed fold changes (mutant/WT). **(A-1)**, Lipid/fatty-acid associated metabolites; **(A-2)**, Nucleotide-related metabolites. Red: relatively high abundance; blue: relatively low abundance. Lipid class abbreviation followed by number of C-atoms: number of double bonds; suffixes “a” and “b” denote each isoform. **(B)** Schematic representation of lipid/fatty acid and nucleotide metabolism in Chlamydomonas. Solid arrows represent single reaction steps, and dashed arrows represent multiple sequential reaction steps. Metabolites shown in blue denote significantly decreased metabolites. Abbreviations: Ac-CoA, acetyl-CoA; Mal-CoA, malonyl-CoA; 3-Oxo-CoA, 3-oxooctanoyl-CoA; MyA, myristic acid; GLA, γ-linolenic acid; EPA, eicosapentaenoic acid; TAG, triacylglycerol; PE, phosphatidylethanolamine; R5P, ribose 5-phosphate; 2dRib, 2-deoxy-D-ribose; dR5P, deoxyribose 5-phosphate; PRPP, phosphoribosyl pyrophosphate; IMP, inosine monophosphate; AMP, adenosine monophosphate; dAMP, 2′-deoxyadenosine monophosphate; ADPR, ADP-ribose; XMP, xanthosine monophosphate; dGMP, 2′-deoxyguanosine monophosphate; cGMP, cyclic GMP; 1,7-DMX, 1,7-dimethylxanthine; Orotidine-5P, orotidine 5′-phosphate; UMP, uridine monophosphate; CMP, cytidine monophosphate; Cyt, cytosine; dTMP, deoxythymidine monophosphate.

In addition to lipids, 13 nucleotide-related DAMs were detected, all reduced in the *tpt10* mutants relative to WT cells (Supplemental Table 1). 2-deoxy-D-ribose and deoxyribose 5-phosphate, associated with deoxyribonucleoside turnover and salvage, decreased to 44–45% and 56–62% of the level in WT cells, respectively (Figures 4A-2 and 4B; Supplemental Figure 6; Supplemental Table 1). Metabolites associated with purine and pyrimidine nucleotide products were also reduced; these metabolites include 7 purine-related and 4 pyrimidine-related products (Figures 4A-2 and 4B; Supplemental Table 1).

Together with the observed reductions in lipid metabolites, these results indicate that the synthesis of primary metabolites, including chloroplast-localized fatty acids and nucleotides, is broadly impaired in the mutants under the tested conditions.

### Changes in Secondary Metabolites *tpt10* Mutants in the Dark

Along with changes in primary metabolism, we observed a marked reduction in the pool of secondary metabolites. 7 isoprenoid-related DAMs were identified, all of which were decreased relative to levels in WT cells (Supplemental Table 1). The 2-C-methyl-D-erythritol 4-phosphate (MEP) pathway–derived prenyl diphosphate precursor dimethylallyl diphosphate declined to 56–57% of that in WT cells (Figures 5A-1 and 5B; Supplemental Figure 6; Supplemental Table 1). Consistent with this upstream change, 6 isoprenoid products showed reduced abundance, including the chloroplast-synthesized carotenoid zeaxanthin, the tocochromanols α-tocopherol and δ-tocotrienol (δ-T3), as well as the plastid-initiated hormones abscisic acid, gibberellin A9, and the cytokinin dihydrozeatin (Figures 5A-1 and 5B; Supplemental Figure 6; Supplemental Table 1).

**Figure 5.**
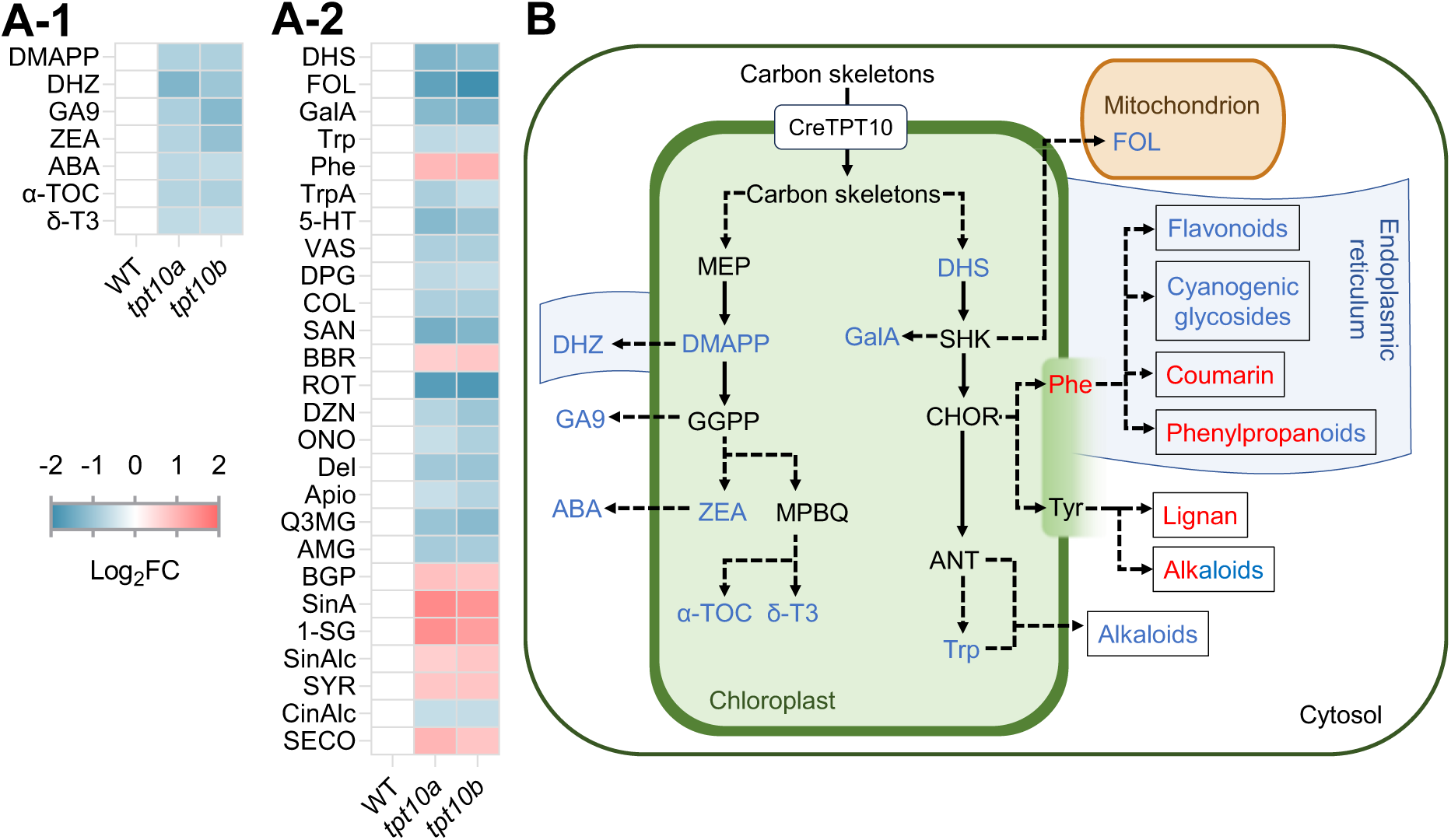
Changes in metabolites associated with plastid-initiated secondary metabolism in WT, *tpt10a*, and *tpt10b* after 48 h of darkness. **(A)** Heatmaps of significantly altered metabolites associated with plastid-initiated secondary metabolism after 48 h of extended darkness, shown as log_2_-transformed fold changes (mutant/WT). **(A-1)**, Isoprenoid-related metabolites; **(A-2)**, Shikimate- and aromatic amino acids-related metabolites. Red: relatively high abundance; blue: relatively low abundance. **(B)** Schematic representation of chloroplast-initiated secondary metabolism in Chlamydomonas. Solid arrows represent single reaction steps, and dashed arrows represent multiple sequential reaction steps. Metabolites shown in red and blue denote significantly increased and decreased metabolites, respectively. Shaded gradient regions denote metabolite groups for which plastid-derived precursors are supplied by the chloroplast. Abbreviations: MEP, 2-C-methyl-D-erythritol 4-phosphate; DMAPP, dimethylallyl diphosphate; DHZ, dihydrozeatin; GGPP, geranylgeranyl diphosphate; GA9, gibberellin A9; ZEA, zeaxanthin; ABA, abscisic acid; MPBQ, methyl-6-phytyl-1,4-benzoquinone; α-TOC, (+)-α-tocopherol; δ-T3, δ-tocotrienol; DHS, 3-dehydroshikimate; SHK, shikimate; GalA, gallic acid; FOL, folic acid; CHOR, chorismate; ANT, anthranilate; Trp, tryptophan; Phe, phenylalanine; Tyr, tyrosine; TrpA, tryptamine; 5-HT, serotonin; VAS, vasicine; DPG, deoxypumiloside; BBR, berberine; COL, colchicine; SAN, sanguinarine; ROT, rotenone; DZN, daidzin; ONO, ononin; Del, delphinidin; Apio, apioside; Q3MG, quercetin 3-O-malonylglucoside; AMG, amygdalin; BGP, bergaptol; SinA, sinapic acid; 1-SG, 1-O-sinapoyl-β-D-glucose; SinAlc, sinapyl alcohol; CinAlc, cinnamyl alcohol; SYR, eleutheroside B; SECO, secoisolariciresinol.

Metabolites associated with the shikimate pathway and downstream reactions involved in the synthesis of aromatic amino acids were also substantially affected, with 26 DAMs identified of which 18 decreased and 8 increased relative to WT cells (Supplemental Table 1). The shikimate intermediate 3-dehydroshikimate declined to 38–42% of WT, accompanied by diminished levels of downstream products including gallic acid and folic acid (Figures 5A-2 and 5B; Supplemental Figure 6; Supplemental Table 1). Furthermore, chloroplast-localized tryptophan decreased in the *tpt10* mutants, while phenylalanine and tyrosine, synthesized in both the chloroplast and cytosol, were elevated relative to levels in WT cells (Figures 5A-2 and 5B; Supplemental Table 1). Finally, 21 secondary metabolites derived from aromatic amino acids showed differential changes, with 14 decreased, which includes alkaloids, flavonoids, a phenylpropanoid, and a cyanogenic glycoside, and 7 increased, including phenylpropanoids, a coumarin, an alkaloid, and a lignan (Figures 5A-2 and 5B; Supplemental Table 1).

In sum, the loss of CreTPT10 broadly disrupted secondary metabolism, with decreases in isoprenoid- and shikimate-derived metabolites, including key intermediates of chloroplast-synthesized products and downstream aromatic amino acids derived from these compounds.

### Transcriptional Changes in the *tpt10* Mutants during Extended Darkness

To investigate whether the metabolic alterations observed in *tpt10* mutants were accompanied by transcriptional changes, we performed RNA-seq during the light-to-dark transition. PCA revealed a pronounced time-dependent progression along PC1 (74.1% of variance) from 0 h to 48 h of darkness, whereas PC2 (14.0%) captured differences between WT cells and *tpt10b* (Supplemental Figure 7A). Biological replicates clustered tightly within each group, indicating high reproducibility. Hierarchical clustering further resolved distinct temporal expression patterns during extended darkness for both WT and the *tpt10* mutant. In WT, transcriptomes shifted progressively over time, while the *tpt10* mutant displayed expression patterns that differed from WT across multiple gene clusters at all time points (Figure 6A). Consistently, the mean Z-score across genes showed a stronger decrease at 24 h followed by a rebound at 48 h in the *tpt10b* mutant, while WT displayed more moderate temporal fluctuations (Figure 6B). Differential expression analysis identified extensive transcriptional changes in the *tpt10b* mutant at both 24 h and 48 h of extended darkness (|log_2_FC| > 1, P-adjust < 0.05; Supplemental Data Set 3). At 24 h, 4,503 genes were differentially expressed relative to WT (1,592 up-regulated; 2,911 down-regulated), whereas by 48 h this number decreased to 837 (225 up-regulated; 612 down-regulated) (Supplemental Figures 7B and 7C; Supplemental Data Set 3). This attenuation of transcriptional divergence at 48 h was also evident in volcano plots, which showed fewer and smaller expression changes compared with 24 h (Supplemental Figures 7B and 7C).

**Figure 6.**
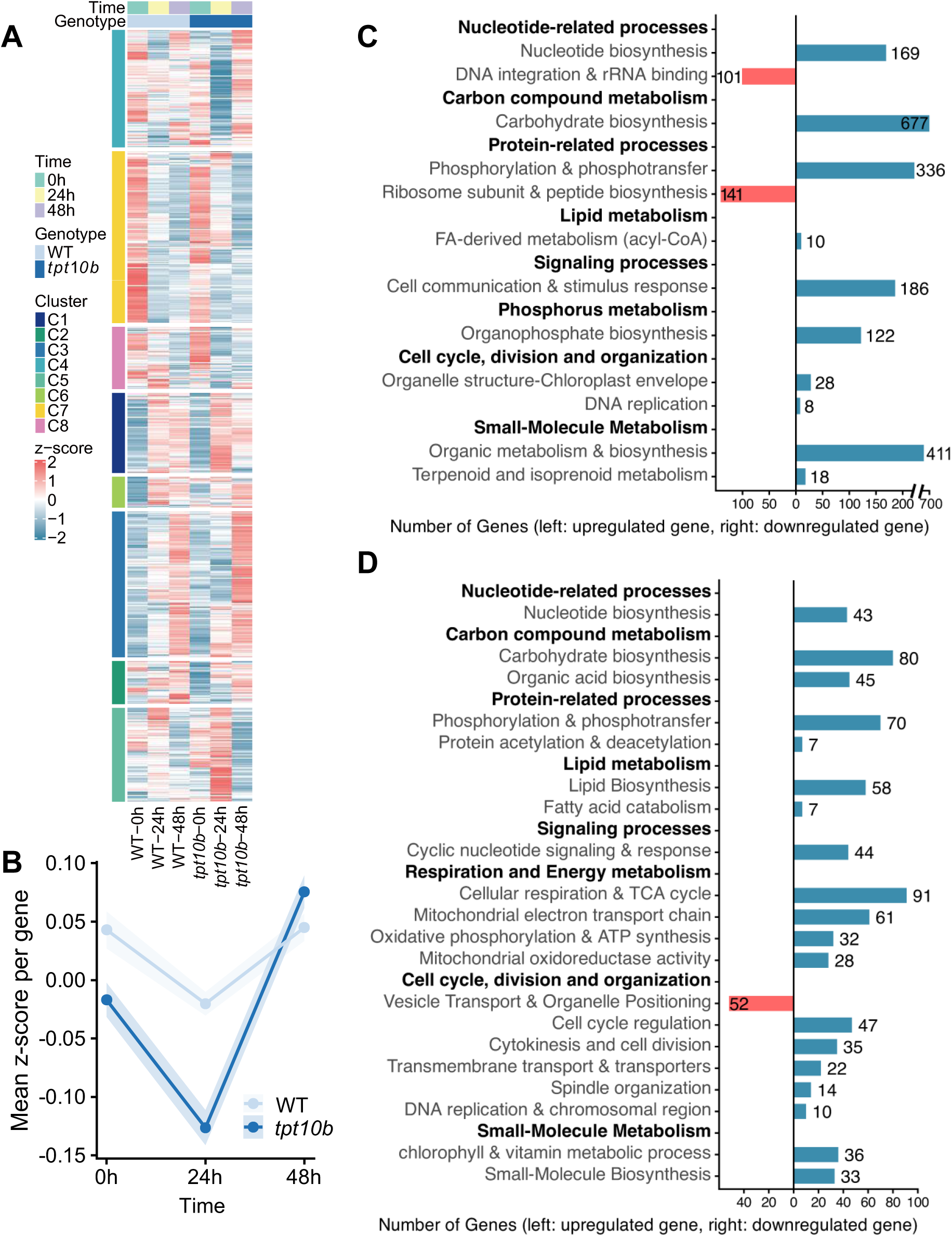
Global transcriptomic reprogramming in the *tpt10b* mutant under extended darkness. **(A)** Hierarchical clustering heatmap of all genes across genotypes (WT and *tpt10b*) and time points (0, 24, and 48 h). Expression values are shown as row-wise Z-scores, calculated for each gene across all samples. **(B)** Temporal dynamics of global gene expression profiles in WT and *tpt10b* under extended darkness, shown as the mean Z-score per gene at each time point. **(C and D)** Functional categorization of significantly enriched GO terms for DEGs at 24 h **(C)** and 48 h **(D)** of darkness (Q < 0.05).

**Figure 7.**
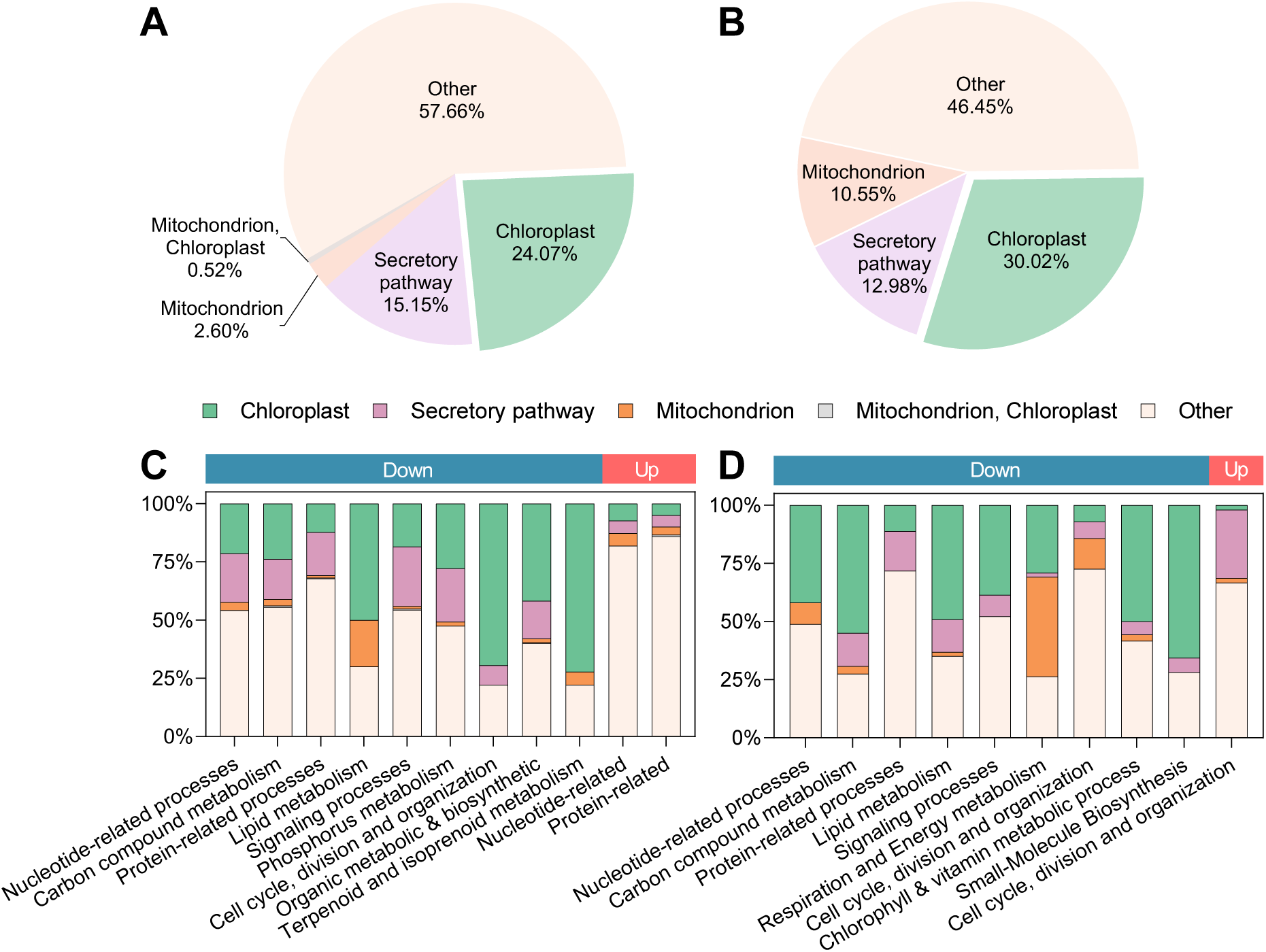
Subcellular localization of GO functional categories enriched in DEGs under darkness. (A and. **B)** Pie charts of PB-Chlamy-predicted subcellular localization of DEGs spanning all GO functional categories for the *tpt10* mutant relative to WT after 24 h **(A)** and 48 h **(B)** of extended darkness. **(C and D)** Stacked bar charts of PB-Chlamy-predicted subcellular localization composition for DEGs within each GO functional category after 24 h **(C)** and 48 h **(D)** of extended darkness. For pie charts, percentages indicate the fraction of DEGs assigned to each PB-Chlamy localization category, including Chloroplast, Secretory pathway, Mitochondrion, dual-targeted (Mitochondrion, Chloroplast), and Other. Stacked bar charts use the same localization categories, and each bar is normalized to 100% within a GO functional category. ‘Other’ denotes genes that could not be assigned to a specific, discernible localization class by PB-Chlamy. The color strips above the stacked bars represent down-regulated genes (blue) and up-regulated genes (red).

Across both dark time points, transcriptional changes in the *tpt10* mutant were dominated by down-regulation of genes associated with biosynthetic and energetic pathways, with several functional modules exhibiting early disruption at 24 h that intensified into broader repression by 48 h (Figures 6C and 6D; Supplemental Table 2; Supplemental Data Set 4). At 24 h, repression was most prominent in carbohydrate biosynthesis, encompassing 677 down-regulated genes, and although fewer genes were affected at 48 h, suppression of this pathway persisted (Figures 6C and 6D; Supplemental Table 2; Supplemental Data Set 4). Similarly, protein-related processes were also suppressed, with 336 down-regulated genes at 24 h and 70 at 48 h (Figures 6C and 6D; Supplemental Table 2; Supplemental Data Set 4). In parallel, nucleotide metabolism was strongly repressed at 24 h, with 169 down-regulated genes assigned to “nucleotide biosynthesis” and this repression persisted at 48 h based on a smaller gene set (43 genes) (Figures 6C and 6D; Supplemental Table 2; Supplemental Data Set 4).

In contrast to these early-responding pathways, fatty acid-related processes displayed a more gradual response. Only 10 genes associated with fatty acid-derived metabolism (acyl-CoA) were repressed at 24 h; however, by 48 h, lipid metabolic suppression had broadened to encompass both fatty acid biosynthesis (58 genes) and catabolism (7 genes) (Figures 6C and 6D; Supplemental Table 2; Supplemental Data Set 4).

Beyond primary metabolism, genes associated with biosynthesis of secondary metabolites were also strongly down-regulated, including those in the MEP-derived isoprenoid pathway (Figure 6C). Isoprenoid biosynthesis via the plastid-localized MEP pathway was already repressed at 24 h, with 18 downregulated genes assigned to genes assigned the to “terpenoid and isoprenoid metabolism” (Figure 6C; Supplemental Table 2; Supplemental Data Set 4). By 48 h, repression extended to the biosynthesis of chlorophyll and carotenoids, encompassing 36 down-regulated genes in “chlorophyll metabolic process” (Figure 6D; Supplemental Table 2; Supplemental Data Set 4).

Collectively, the loss of CreTPT10 led to widespread repression of biosynthetic pathways in primary and secondary metabolism, many of which showed early suppression that persisted or expanded during extended darkness.

### Transcriptional Changes Associated with Energy Metabolism and Regulatory Pathways in the *tpt10* Mutant during Extended Darkness

Notably, a distinct temporal pattern was also observed for mitochondrial functions. Genes involved in mitochondrial energy metabolism were largely repressed, with this effect was evident only after 48 h in darkness (Figure 6D). At this time point, over 200 genes associated with aerobic respiration and the mitochondrial electron transport chain were down-regulated across core energy-production pathways (Figure 6D; Supplemental Table 2; Supplemental Data Set 4). This coordinated transcriptional repression corresponded to the observed decline in respiration rates in *tpt10* strains, indicating impaired mitochondrial respiratory capacity under extended darkness. Signaling- and cell cycle-related processes were also affected during extended darkness. At 24 h, signaling pathways were strongly impacted, with 186 down-regulated genes being annotated as associated with cell communication and stimulus response processes (Figure 6C; Supplemental Table 2; Supplemental Data Set 4). At this time point, changes in cell cycle-related functions were limited, whereas 28 down-regulated genes were linked to chloroplast envelope organization (Figure 6C; Supplemental Table 2; Supplemental Data Set 4). At 48 h, transcriptional changes were detected across cell cycle, division, and organization-related categories, involving more than 100 genes in total, with the majority showing reduced expression levels (Figure 6D; Supplemental Table 2; Supplemental Data Set 4). Signaling processes were also affected at this time point, with 44 down-regulated genes assigned to cyclic nucleotide signaling and response pathways (Figure 6D; Supplemental Table 2; Supplemental Data Set 4).

Overall, beyond biosynthetic metabolism, loss of CreTPT10 perturbed mitochondrial and cellular regulatory programs, highlighting its central role in sustaining anabolic and energetic metabolism under extended darkness.

### Compartmentalized Distribution of Transcriptional Reprogramming in the *tpt10* Mutant

To further understand the spatial context of transcriptional reprogramming under extended darkness, the subcellular localization of proteins encoded by DEGs associated with enriched GO terms was predicted using TargetP and PB-Chlamy, which were highly concordant (Figures 7A and 7B; Supplemental Figures 8A and 8B; Supplemental Table 2). Globally, aside from DEGs encoding proteins without clear localization signals, genes encoding proteins predicted to localize to the chloroplast predominated at both 24 h (24.1%) and 48 h (30.0%), whereas genes encoding proteins predicted to localize to mitochondria were initially scarce at 24 h (2.6%) but increased by 48 h (10.6%) (Figures 7A and 7B; Supplemental Table 2).

Within each GO category, genes encoding chloroplast-localized proteins dominated most biosynthetic and metabolic processes, including lipid, terpenoid/isoprenoid, and carbon compound metabolism (typically 40–70%; Figures 6C and 6D; Supplemental Table 2), with additional contributions from secretory pathway-localized proteins (15–25%; Figures 7C and 7D; Supplemental Table 2). By 48 h, genes encoding mitochondrial-localized proteins were selectively enriched in respiration and energy metabolism (42.7%) (Figures 7D; Supplemental Table 2). Transcriptional repression in the *tpt10* mutant in the dark is spatially organized, with repression of chloroplast-associated genes being the most dominant relative to the other metabolic genes that are repressed. However, nuclear genes encoding mitochondrial proteins are also repressed in the mutant, with those associated with energy metabolism enriched at the later time point.

## Discussion

### CreTPT10 Functions as a Major Transporter for X5P Across the Chloroplast Envelope

In the yeast liposome assay, CreTPT10 exhibited a strong preference for both X5P and dihydroxyacetone phosphate, with the highest transport activity observed for X5P (Figure 3), indicating that these metabolites represent potential physiological substrates of CreTPT10 in vivo. However, multiple independent lines of evidence converge on X5P as the primary physiological substrate of CreTPT10 in vivo. In Chlamydomonas, CreTPT2 and CreTPT3 have been reported to specifically transport dihydroxyacetone phosphate, a triose phosphate (Huang et al., 2023). To assess whether triose phosphate transport contributes to heterotrophic growth, we examined the growth performance of *tpt2* and *tpt3* mutants under dark conditions. Notably, both mutants displayed enhanced growth relative to the WT (Supplemental Figure 9), indicating that loss of dihydroxyacetone phosphate transport does not compromise, and may even promote, heterotrophic growth in the dark. In contrast, loss of CreTPT10 resulted in a pronounced growth inhibition under the same conditions, suggesting that X5P, rather than dihydroxyacetone phosphate, represents the major physiological substrate supporting CreTPT10-dependent plastid metabolism in darkness. Consistent with this interpretation, *tpt10* mutants showed a marked accumulation of X5P, compared with the WT (Figure 8; Supplemental Figure 10), implying that plastid utilization of X5P is impaired upon loss of CreTPT10. Taken together, CreTPT10 preferentially transports X5P across the chloroplast envelope to sustain cellular growth under heterotrophic conditions.

**Figure 8.**
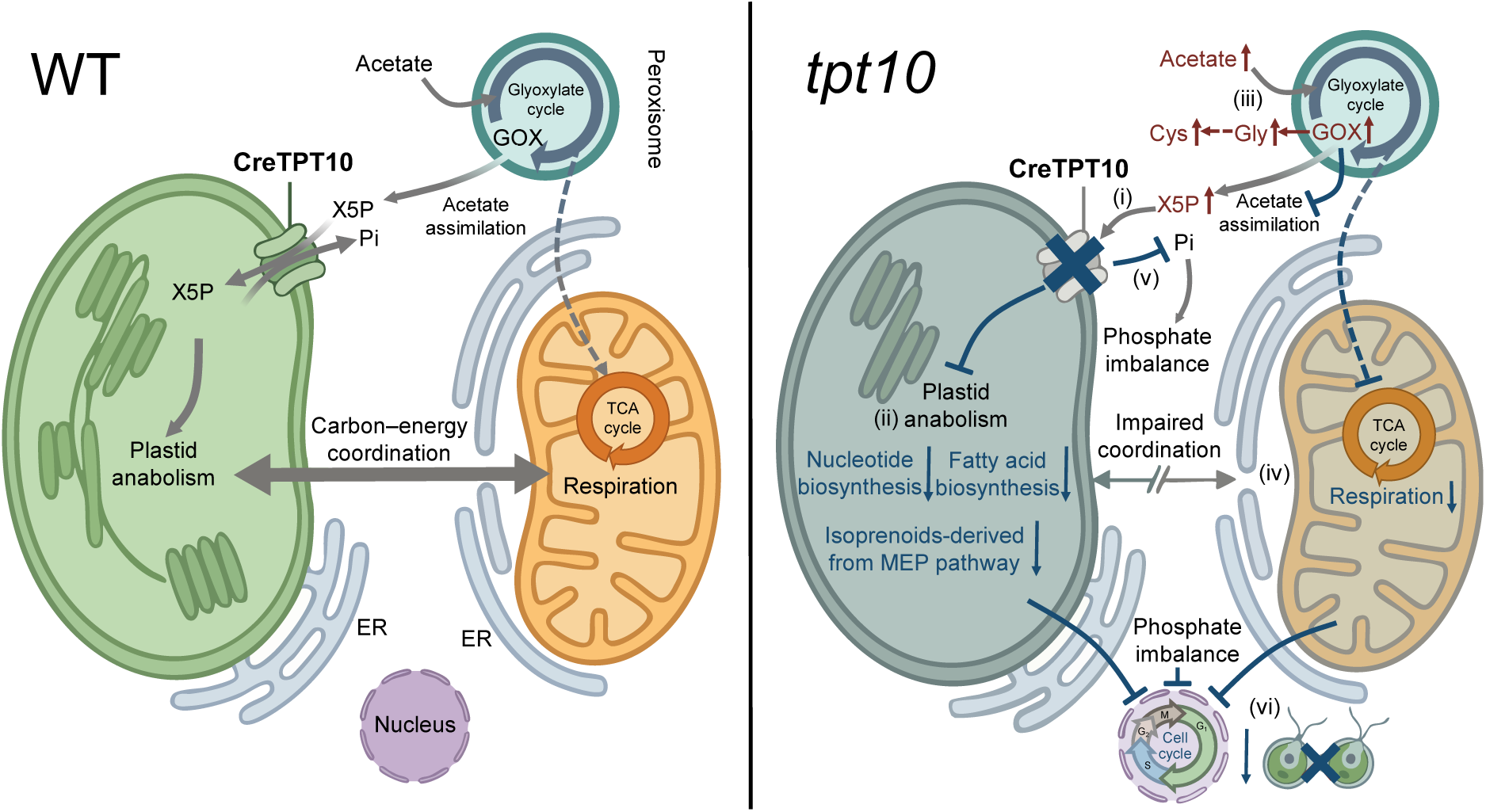
Effect of eliminating Chlamydomonas CreTPT10 on carbon allocation and organelle metabolic coordination in the dark. Schematic model comparing WT (left) and tpt10 mutant (right) during dark heterotrophic growth on acetate. **(i)** Loss of CreTPT10 impairs chloroplast import of cytosolic X5P, leading to X5P accumulation. **(ii)** Reduced X5P import is accompanied by collapse of chloroplast anabolic metabolism. **(iii)** Cytosolic X5P accumulation creates a metabolic bottleneck in gluconeogenesis and the glyoxylate shunt, leading to accumulation of pathway intermediates, diversion of carbon into Gly and Cys biosynthesis, and elevated acetate levels. **(iv)** Mitochondrial respiratory capacity is diminished. **(v)** Disrupted X5P/Pi exchange perturbs phosphate homeostasis. **(vi)** The combined impairment of plastid anabolism, respiration, and phosphate-dependent metabolism constrains cell-cycle progression and division. Abbreviations: X5P, xylulose 5-phosphate; GOX, glyoxylate; Gly, glycine; Cys, cysteine; Pi, inorganic phosphate; TCA, tricarboxylic acid.

### CreTPT10 Imports Carbon Skeletons into the Chloroplast to Sustain Anabolism

Loss of CreTPT10 impaired growth in darkness and triggered extensive transcriptomic and metabolomic reprogramming relative to the WT (Figures 4, 5 and 6; Supplemental Tables 1 and 2). In contrast, single knockout of *Arabidopsis* XPT displays no obvious visible phenotype, underscoring the greater dependence of Chlamydomonas plastid metabolism on XPT-mediated carbon import when photosynthetic carbon fixation is absent. Furthermore, loss of CreTPT10 constrained plastid-localized anabolic metabolism, reflecting a coordinated reduction in plastid anabolic capacity rather than isolated pathway defects during extended darkness (Figures 4 and 5; Supplemental Table 1). Notably, in liquid culture, *tpt10* mutants ultimately reached cell densities comparable to WT at stationary phase despite pronounced early growth defects, suggesting that additional transporters can partially compensate for organic carbon delivery into the chloroplast.

In Chlamydomonas, de novo fatty acid synthesis occurs in the chloroplast, and plastid-derived acyl supply supports lipid production (Fan et al., 2011; Boyle et al., 2012; Li-Beisson et al., 2015; Warakanont et al., 2015; Choi et al., 2022). In *tpt10* mutants, acetyl-derived precursors were reduced, accompanied by a broad decrease in the fatty acids and lipid pools (Figure 4A-1; Supplemental Table 1), consistent with impaired plastidial provision of carbon skeletons for de novo fatty acid synthesis. GO enrichment analysis of downregulated genes further highlighted reduced lipid biosynthesis, with the majority encoding proteins predicted to localize to the chloroplast (Figures 6C, 6D, 7C and 7D; Supplemental Table 2). Transcriptome profiling also showed broad downregulation of carbohydrate biosynthesis and organic acid biosynthesis, and the associated genes largely encoded chloroplast-targeted proteins.

Similarly, de novo nucleotide synthesis in photosynthetic eukaryotes is compartmentalized: purine biosynthesis occurs mainly in plastids, whereas pyrimidine synthesis initiates in plastids before continuing in the mitochondria and cytosol (Doremus and Jagendorf, 1985; Hung et al., 2004; Zrenner et al., 2006; Witz et al., 2012; Witte and Herde, 2020). In *tpt10* mutants, metabolomics revealed reductions in the nucleotide pool, with reduced ribose-derived intermediates and broad decreases across both purine- and pyrimidine-associated metabolites relative to WT cells (Figure 5A-2; Supplemental Table 1). Correspondingly, GO enrichment analysis of downregulated genes indicated decreased nucleotide biosynthesis, with the majority of these genes encoding chloroplast-targeted proteins (Figures 6C, 6D, 7C and 7D; Supplemental Table 2).

Moreover, isoprenoid precursors in Chlamydomonas are exclusively generated via the plastidial MEP pathway, positioning the chloroplast as the central hub for isoprenoid precursor supply (DellaPenna and Pogson, 2006; Vranová et al., 2013; Zhao et al., 2022; Lanier et al., 2023). Metabolomics showed reduced isoprenoid precursor pool in *tpt10* mutants, accompanied by decreases in plastid-synthesized isoprenoid products such as carotenoids and tocochromanols (Figure 5A-1; Supplemental Table 1), as well as lower isoprenoid hormones whose downstream conversions occur outside the chloroplast (Helliwell et al., 2001; Cheng et al., 2002; Takei et al., 2004; Shumskaya et al., 2012; Romer et al., 2024; Sakakibara, 2025; Figure 5A-1; Supplemental Table 1). Transcriptome profiling revealed repression of terpenoid and isoprenoid metabolism, with a clear bias toward chloroplast-targeted proteins (Figures 6C and 7C; Supplemental Table 2).

Beyond the MEP pathway, the plastid-localized shikimate pathway provides another major precursor for aromatic amino acid biosynthesis and secondary metabolism (Schmid and Amrhein, 1995; Herrmann and Weaver, 1999; Pott et al., 2019). Metabolomics showed reduced levels of an upstream shikimate intermediate, together with coordinated decreases in plastid-derived aromatic amino acids and a subset of plastid-initiated secondary metabolites in *tpt10* mutants (Figure 5A-1; Supplemental Table 1). Although increases were detected among selected downstream aromatic secondary metabolites, these changes did not converge on a single branch.

Finally, loss of CreTPT10 triggered metabolic alterations that were embedded within a broader pattern of coordinated transcript–metabolite variation. Gene-metabolite correlation analysis revealed coordinated transcript-metabolite variation, with strong positive and negative associations (Supplemental Figure 11A). Within this framework, 4-diphosphocytidyl-2C-methyl-D-erythritol synthase (CMS1) emerged as a highly connected chloroplast node linked to multiple metabolite classes, indicating that MEP pathway transcription tracks global metabolic state rather than functioning as an isolated regulatory driver and highlighting that CreTPT10-mediated carbon import influences this coordinated plastid metabolic network (Supplemental Figure 11B and 11C).

Together, these results demonstrate that chloroplast-localized anabolic metabolism depends on X5P imported by CreTPT10 for carbon skeleton supply and coordination of plastid metabolic networks during extended darkness.

### CreTPT10 Supports Optimal Energy Coordination between the Chloroplast and Mitochondria

Loss of CreTPT10 impaired chloroplast import and likely limited the utilization of cytosolic X5P, resulting in its accumulation in the cytosol (Figure 8), which was indeed higher in *tpt10* mutants than in WT cells in the dark (Supplemental Figure 10; Supplemental Table 1). Consistent with this defect, failure to import X5P was accompanied by a marked collapse of chloroplast anabolic metabolism (Figure 8).

In addition, cytosolic X5P buildup may constrain upstream carbon flow into gluconeogenesis, thereby creating a metabolic bottleneck that feeds back to hinder the glyoxylate shunt (Figure 8). Supporting this model, gluconeogenic intermediates (phosphoenolpyruvate and glyceraldehyde 3-phosphate) and glyoxylate-cycle metabolites (glyoxylate, succinate, and malate) accumulated in *tpt10* mutants relative to WT (Supplemental Figure 10; Supplemental Table 1). Part of this excess carbon was redirected into amino acid biosynthesis (glycine and cysteine), suggesting an overflow outlet for the stalled pathway (Supplemental Figure 10; Supplemental Table 1). Such metabolic restriction may further impair acetate assimilation, as indicated by elevated acetate levels in *tpt10* cells (Supplemental Figure 10; Supplemental Table 1).

Collectively, disruption of CreTPT10 dampened chloroplast anabolic activity and hindered carbon flux through gluconeogenesis and the glyoxylate shunt in the dark, while also reducing acetate utilization. These changes together limited substrate availability for mitochondrial respiration, leading to decreased respiratory capacity and broad repression of nuclear genes encoding mitochondrial electron transport chain components (Figure 6D; Supplemental Table 2). Because XPT mediates counter-exchange of inorganic phosphate (Pi) and X5P across the chloroplast envelope, loss of CreTPT10 likely also perturbs phosphate homeostasis (Figure 3). Consistent with this possibility, transcriptomic analysis revealed widespread repression of organophosphate biosynthesis and phosphorylation-related processes (Figure 6C; Supplemental Table 2). Ultimately, impairment of these interconnected anabolic and energetic pathways constrained cell proliferation, as evidenced by reduced growth and extensive downregulation of cell cycle-associated genes in the *tpt10* mutant (Figure 6D; Supplemental Table 2).

Collectively, our findings position CreTPT10 as an integrative hub that couples chloroplast anabolic activity with mitochondrial respiration, phosphate homeostasis, and acetate assimilation to maintain proliferative capacity during prolonged darkness.

In conclusion, we identified CreTPT10 as a chloroplast envelope transporter that mediates X5P import from the cytosol. Our results demonstrate that importing X5P provides the carbon skeleton for chloroplast anabolism, including lipids, nucleotides, and isoprenoids. Furthermore, the loss of CreTPT10 is associated with a respiratory downshift and disruption of carbon-energy coordination between the chloroplast and mitochondria. Overall, our findings expand the functional scope of plastid phosphate translocators, revealing that pPTs are not restricted to exporting carbon during photosynthesis but also facilitate carbon import to sustain growth under heterotrophic conditions.

### Outlook

For decades, studies of plastid phosphate translocators have focused primarily on their roles in photosynthetic carbon export and source-sink partitioning in higher plants, with comparatively little attention to their functions under non-photosynthetic or heterotrophic conditions. How plastids in unicellular algae sustain anabolic metabolism in the absence of photosynthetic carbon fixation has therefore remained poorly understood. Here, we show that CreTPT10 imports X5P into the Chlamydomonas chloroplast, providing carbon skeletons for anabolic metabolism and facilitating inter-organelle energy interactions, thereby supporting compartmentalized carbon allocation and cellular metabolic homeostasis. Our future work will elucidate how plastids maintain anabolic metabolism under heterotrophic or dark conditions through the coordinated action of additional transporters, and to define compensatory pathways activated upon loss of CreTPT10, as well as their contribution to restoring plastid function and cellular metabolic balance. Addressing these questions will be critical for understanding how plastid carbon import is dynamically regulated in algae and contributes to metabolic flexibility under multiple trophic conditions.

## Methods

### Strains and Culture Conditions

The WT Chlamydomonas, strain M10 (CC-4403), an isogenic derivative of CC-124, was used as the parental strain to generate knockout mutants. Algal cultures were routinely cultivated in growth chambers at 25 °C with continuous shaking (150 rpm) in Tris-Acetate-Phosphate (TAP) medium. Cultures were illuminated with continuous cool white LEDs at low light (LL, 30 μmol photons m^-2^ s^-1^). For growth assays, cultures were inoculated at a starting density of 5 × 10^5^ cells mL^−1^ in TAP with shaking in the dark. Chlorophyll concentrations were measured by spectrophotometry according to Porra et al. (1989). For cell density determinations, 200 μL of cell suspension was fixed with 3 μL of Lugol’s iodine solution, and the cell density quantified using a hemocytometer. Three biological replicates were used to assay the samples. Growth assays on solid medium were performed with cultures spotted onto TAP-agar (1.0 agar) at various chlorophyll concentrations (4, 2, 1, 0.5 μg mL^-1^) and exposed to different light conditions: continuous light (LL, or high light, HL, 50 μmol photons m^-2^ s^-1^), diurnal cycle (DC, 12 h D/12 h L, 30 μmol photons m^-2^ s^-1^), and extended darkness. In addition, growth on agar plates under nitrogen-deficient conditions (−N, 30 μmol photons m^-2^ s^-1^, continuous light) was tested. These agar plates were incubated for 24 days at 25 °C.

### Vector Construction, Transformation, and Subcellular Localization

The pRam118_*VENUS* plasmid, harboring the *VENUS* gene and the *AphVII* cassette (conferring resistance to hygromycin), was used to construct the *CreTPT10* (Cre08.g379350_4532) expression vector (Kaye et al., 2019). The pRam118_VENUS backbone was amplified by PCR as a linear plasmid using the primers pRam118_f and pRam118_r (Supplemental Table 3). The full-length *CreTPT10* coding sequence (CDS) was synthesized by Tsingke Biotech (Beijing, China). The primers *TPT10*CDS_f and *TPT10*CDS_r (Supplemental Table 3) were used to amplify *CreTPT10* CDS fragments containing an overlap with the linearized pRam118_VENUS vector. For generating the plasmid pRam118_*CreTPT10*&*VENUS*, full-length *CreTPT10* CDS was assembled with the pRam118_VENUS plasmid using Gibson assembly (Gibson et al., 2009). The resulting construct was designated pRam118_*CreTPT10*&*VENUS*, which includes the *PsaD* promoter, the full-length *CreTPT10* CDS, *VENUS*, the *AphVII* cassette, and the *RBCS2* 3’UTR (Supplemental Figure 12).

For each transformation, 2–4 μg of linearized plasmid was added to 250 μL of a cell suspension at ∼2 × 10^8^ cells mL^-1^. GeneArt MAX Efficiency Transformation Reagent for algae (Invitrogen, Carlsbad, CA, USA) was used for electroporation according to the manufacturer’s instructions. Electroporation was performed using the GenePulser II electroporator (Bio-Rad, Hercules, CA, U A) at 500 V, 50 μF, and 800 Ω. Transformants were selected on TAP agar medium containing 10 μg mL^-1^ hygromycin. Drug-resistant transformants were screened and visualized for VENUS fluorescence as described previously (Huang et al., 2023). In brief, fluorescence was initially examined using a microplate reader (Molecular Devices SpectraMax i3x, Sunnyvale, CA, USA). The VENUS fluorophore was excited at 515 nm and fluorescence emission screened at 550 nm (bandwidths of 12 nm). Chlorophyll autofluorescence was assessed by exciting the cells at 440 nm and monitoring emission at 680 nm (bandwidths of 9 nm and 20 nm, respectively). High-resolution fluorescence imaging was performed with a TCS SP8 confocal laser-scanning microscope (Leica Microsystems, Wetzlar, Germany). Excitation and emission settings for the microscope were as follows: VENUS fluorescence excitation was at 514 nm and the emission detected between 525 and 575 nm using a HyD SMD hybrid detector, whereas chlorophyll autofluorescence was excited at 488 nm and emission detected between 680 and 720 nm using a HyD SMD hybrid detector (Blaby-Haas et al., 2018).

### CRISPR/Cas9-Mediated Mutagenesis and RT-qPCR Verification

WT cells were cultured under continuous LL for 3 d to a density of 3 to 5 × 10^6^ cells mL^-1^, and then concentrated to 2 × 10^8^ cells mL^−1^ in 0.5 × TAP medium supplemented with 40 mM sucrose to enhance electroporation survival. A single-guide RNA (sgRNA) targeting exon 9 of *CreTPT10* was designed using CHOPCHOP (https://chopchop.cbu.uib.no/), yielding the sequence Cre*TPT10*-sg (5′-GUUGCGGAAGAACAGCACCGAGG-3′). Exon 9 was selected because it encodes part of the conserved transporter core, and frameshift mutations at this site are expected to abolish function by truncating essential transmembrane segments. The protocol for *CreTPT10* disruption was adapted from Huang et al. (2023) and Findinier (2023). Prior to electroporation, Cas9 (Integrated DNA Technologies, Coralville, IA, USA, Cat. no: 1081058) and sgRNA were incubated at 37 °C for 30 min to assemble a ribonucleoprotein (RNP) complex. Approximately 500 ng PCR-amplified *AphVII* cassette DNA was added to the RNP mixture. A 250 μL aliquot was electroporated using the Super Electroporator NEPA21 type II (Nepa Gene, Chiba, Japan). After 16 h recovery in TAP medium containing 40 mM sucrose under LL, cells were plated onto solid TAP medium containing 10 μg mL^−1^ hygromycin. Sense-oriented *AphVII* knock-in lines were identified by PCR using primer pairs in which one primer annealed to the genomic sequence and the other to the *AphVII* insert (Supplemental Table 3). The amplified fragments were sequenced to verify the insertion sites (Hecegene Technology, Wuhan, China).

RT-qPCR was performed as described previously (Huang et al., 2023). Cells were cultured under LL and then transferred to extended darkness for 24 h before harvest. Briefly, total RNA was extracted using the Eastep Super Total RNA Extraction Kit (Promega, Madison, WI, USA, Cat. no: LS1040) and treated with DNase I. First-strand cDNA was synthesized from 1 μg total RNA using the Toloscript ALL-in-one RT EasyMix for qPCR (Tolobio, Shanghai, China, Cat. no: 22107). RT-qPCR was carried out using PerfectStart Visual Green qPCR SuperMix (Transgene, Beijing, China, Cat. no: AQ6201) on CFX96 Touch Real-Time PCR Detection System (Bio-Rad, Hercules, CA, USA). Primers used to quantify *CreTPT10* transcript levels are listed in Supplemental Table 3. Relative mRNA levels were calculated using the ΔC_t_ method (Silver et al., 2006), with the Chlamydomonas β subunit-like polypeptide (*CBLP*) gene serving as the internal control (Supplemental Table 3).

### Measurement of Oxygen Consumption Rate

Respiration rates were measured with a Chlorolab-2 oxygen electrode (Hansatech Instruments, Norfolk, England) following the manufacturer’s instructions. The electrode was calibrated using O_2_-saturated water and by depleting oxygen in the electrode chamber with the addition of excess sodium dithionite. 1.5 mL of Chlamydomonas cell suspension (4–6 μg chlorophyll mL^−1^) was added to the respiration chamber, and oxygen consumption was measured in the dark at 25 °C. The respiration rate was recorded continuously for about 8 min.

### Production of CreTPT10 in Yeast, Reconstitution into Liposomes, and Transport Assays

The N-terminal transit peptide of CreTPT10 was predicted using TargetP 2.0 (Almagro Armenteros et al., 2019), and the sequence encoding the first 63 amino acids was removed to obtain the mature coding sequence. This truncated sequence was codon-optimized for *Saccharomyces cerevisiae* and synthesized (Thermo Fisher Scientific). The insert was cloned into pYES-NTa via BamHI and XhoI, generating an N-terminal His-tag fusion. The yeast expression construct was transformed into *S.cerevisiae* strain INVSc1 using a standard transformation protocol (Gietz and Schiestl, 2007). Expression of the His-tagged mature *CreTPT10* protein was induced by cultivation in galactose-containing medium under control of the GAL1 promoter. Total membranes were isolated from transformed yeast cells and reconstituted into liposomes containing 3% (w/v) phosphatidylcholine. Inorganic phosphate (Pi) transport was measured using radiolabeled orthophosphate (^32^P-Pi; 0.25 mM), as previously described in Huang et al. (2023). As pPT family members function as antiporters, uptake assays were performed in the presence or absence of a 25 mM counter-exchange substrate. Time-course experiments, testing potential substrates, and K_M_/K_i_ analyses were conducted to determine transport kinetics, substrate specificity, and substrate affinities Huang et al. (2023).

### Metabolite Extraction and Metabolic Analysis

Exponential cells grown under continuous LL were transferred to darkness and collected after 4 and 48 h of dark treatment. For each group, a 50 mL culture was rapidly filtered in the dark and quenched in liquid nitrogen (n = 4). Pellets were extracted with 2 mL methanol/acetonitrile/water (2:2:1, v/v/v) using three vortex–freeze–thaw cycles in which the cells were frozen in liquid nitrogen followed by immersion in a 3 °C water bath. After incubation at -20 °C for 1 h, samples were centrifuged at 16,229 × g for 15 min at 4 °C. upernatants were dried in a peedVac Concentrator (Eppendorf, Hamburg, Germany), resuspended in 100 μL water/acetonitrile (1:1, v/v), sonicated at 4 °C for 10 min, centrifuged, and subsequently subjected to analysis by liquid chromatography-mass spectrometry (LC–MS). Metabolites were separated on a BEH Z-HILIC VanGuard FIT column (1. μm, 2.1 × 100 mm, Waters, Milford, MA, U A) using a UHPLC system (12 0 series, Agilent Technologies, Santa Clara, CA, USA) coupled to a quadrupole time-of-flight (Q-TOF) mass spectrometer (6545B Q-TOF, Agilent Technologies, Santa Clara, CA, USA). Samples (4 μL) were injected at a flow rate of 0.4 mL min^-1^ and temperature of 35 °C using both positive and negative electrospray ionization (ESI) modes. The mobile phase A consisted of 25 mM ammonium acetate and 25 mM ammonium hydroxide in water, and the mobile phase B was 100% acetonitrile. The elution gradient for phase B was 78% (0– min), 5 ( –8 min), 5 (8–8.5 min), and 5 (8.5–12 min). Data were acquired using *MassHunter Qualitative Analysis* (v10.0). All solvents and reagents were of chromatographic grade.

LC–MS raw data were preprocessed in *Progenesis QI* (Waters Corporation, Milford, USA) to generate a three-dimensional matrix comprising sample information, metabolite identities, and mass spectrometric response intensities. Artifacts (noise, column bleed, derivatization peaks) were removed, followed by de-replication and peak alignment. Metabolites were identified based on mass-to-charge ratio (m/z), retention time (min), and fragment ion characteristics using the Human Metabolome Database (HMDB, http://www.hmdb.ca/) (Wishart et al., 2022), Metlin (https://metlin.scripps.edu/) (Guijas et al., 2018), and the Meiji Database (MJDB, https://www.majorbio.com/) (Han et al., 2024). Metabolic features detected in at least 80% of the samples in any group were retained. Missing values were imputed using the mean intensity for each metabolite across all samples, and relative metabolite abundances were normalized to cell number. To ensure data quality, we retained only variables that were consistent across quality control (QC) samples, with a relative standard deviation (RSD) of <30%. Pooled QC samples were generated by mixing equal volumes of all sample extracts and were injected periodically during the run to evaluate instrument stability and data reproducibility. Multivariate analysis, including Principal Component Analysis (PCA) and Orthogonal Partial Least Squares Discriminant Analysis (OPLS-DA), was performed using R packages (ropls, v1.6.2) with seven-fold cross-validation (Han et al., 2024). The metabolites with variable importance in projection (VIP) score > 1, p < 0.05 (Student’s t-test) and fold changes (FC) > 1.5 or FC < 0.667 were considered to be differentially abundant metabolites (DAMs) (Sussulini, 2017). DAMs were mapped to KEGG pathways (KEGG, http://www.genome.jp/kegg/), with pathway schematics manually compiled from KEGG reference maps and functional classes assigned using the criteria described by Johnson and Alric (2013).

### RNA Extraction and Transcriptomic Analysis

Following the transition from LL to dark, Chlamydomonas cultures were collected after 0, 24, and 48 h. For each time point, 15 mL of culture from each of three biological replicates was harvested by centrifugation at 3,152 × g and 4 °C for 5 min. The supernatant was discarded, and the cell pellets were immediately snap-frozen in liquid nitrogen and stored at −80 °C until RNA extraction. Total RNA was extracted using the RNAprep Pure Plant Plus Kit (TIANGEN, Beijing, China, Cat. no: DP441) and treated with RNase-Free DNase I. The assessment of total RNA quality and quantity was measured with the NanoDrop 2000 (Thermo Fisher Scientific, Waltham, MA, USA) and Agilent 2100 Bioanalyzer (Agilent Technologies, Santa Clara, CA, USA). Preparation of complementary DNA (cDNA) libraries and transcriptome sequencing were performed at Scale Biomedicine Technology Co., Ltd. (Beijing, China).

Filtered, high-quality reads were aligned to the Chlamydomonas reference genome v6.1 (https://phytozome-next.jgi.doe.gov/info/CreinhardtiiCC_4532_v6_1) (Craig et al., 2023) using Hisat2 (v2.0.5) (Kim et al., 2015). Differential gene expression analyses were performed using the DESeq2 R package (v1.20.0) (Love et al., 2014). Differential expression was determined using adjusted P-value (Benjamini-Hochberg) threshold of P ≤ 0.05 and |log_2_FoldChange (FC)| > 1. Gene Ontology (GO) enrichment was performed using the R package ClusterProfiler (v3.8.1), and terms with Q-values < 0.05 were considered significantly enriched (Ashburner et al., 2000). Protein sequences encoded by DEGs within enriched GO functional categories after 24 h and 48 h of darkness were retrieved from the Chlamydomonas CC-4532 v6.1 genome annotation. The corresponding protein sequences were used as input for TargetP 2.0 to predict subcellular targeting (Almagro Armenteros et al., 2019, https://services.healthtech.dtu.dk/services/TargetP-2.0). PB-Chlamy provides proteome-wide subcellular localization predictions for Chlamydomonas (Wang et al., 2023, https://huggingface.co/wpatena/PB-Chlamy/tree/main). Using the same DEG set within the enriched GO functional categories as in the TargetP analysis, these gene identifiers were matched to the published PB-Chlamy database to retrieve the corresponding predicted subcellular localizations.

### Construction of Phylogenetic Tree

Phylogenetic analyses of 46 pPT sequences from Chlamydomonas, 10 dicot species, and 3 monocot species were performed in MEGA12 using the Maximum Likelihood method with branch lengths in substitutions per site (Yang, 1994; Kumar et al., 2024). The LG + F model with discrete gamma rate variation (+ G; five categories; parameter = 0.7603) was used, branch support was evaluated with 1,000 bootstrap replicates, and trees were visualized in iTOL (https://itol.embl.de/tree).

## Data Availability

RNA-seq data have been deposited in the NCBI Gene Expression Omnibus as GSE320297. Metabolomic data have been deposited at https://ngdc.cncb.ac.cn/omix/ as OMIX014631.

## Author Contributions

W.H. and A.G. conceptualized the study. W.H. generated the CRISPR mutants as well as tested the growth under various conditions. A.W. localized the CreTPT10 proteins, conducted measurements of respiratory O_2_ consumption, extracted samples for LC–MS, performed LC–MS, and carried out metabolomics data analysis. A.W., S.X., and Q.K. performed growth assays. L.S. and N.L. performed the reconstitution into liposomes and transport activity assays. Y.W. and A.W. performed transcriptome analysis. W.H., Y.W., and A.W. wrote the manuscript. A.G., N.L., Y.B. and all the other authors contributed to writing and revising the manuscript.

## Acknowledgements

W.H. thanks Jun Men and Siyu Wang (Analytical Testing Center, Institute of Hydrobiology, Chinese Academy of Sciences) for assistance with metabolite determinations. The authors declare no competing interests.

## Supplemental Information

**Supplemental Figure 1.**
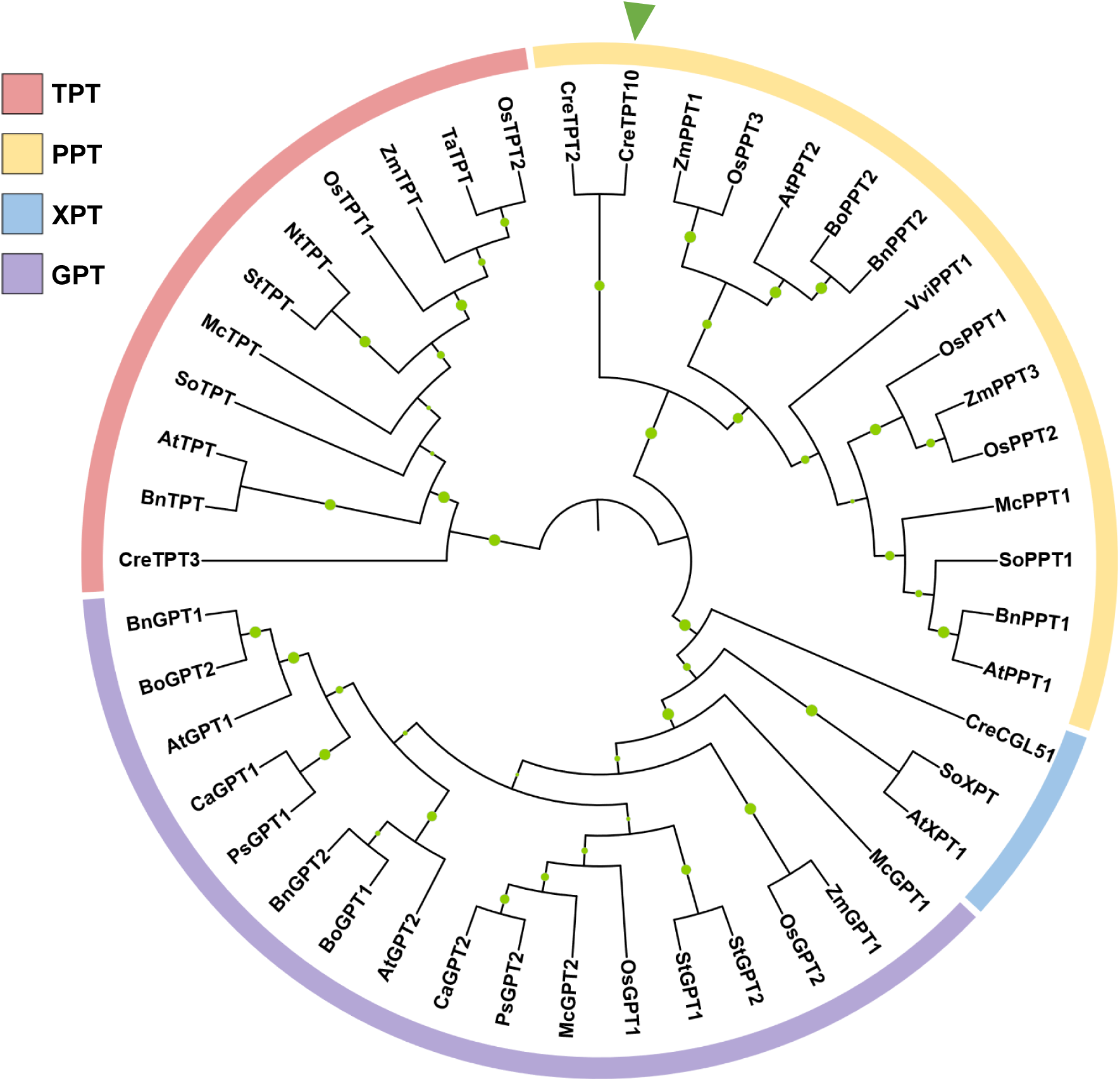
Maximum-likelihood phylogeny of plastidic phosphate translocators (pPTs). Species: *Arabidopsis thaliana* (At), *Brassica napus* (Bn), *Brassica oleracea* (Bo), *Cicer arietinum* (Ca), *Chlamydomonas reinhardtii* (Cre), *Mesembryanthemum crystallinum* (Mc), *Nicotiana tabacum* (Nt), *Oryza sativa* (Os), *Pisum sativum* (Ps), *Spinacia oleracea* (So), *Solanum tuberosum* (St), *Triticum aestivum* (Ta), *Vitis vinifera* (Vvi), *Zea mays* (Zm).

**Supplemental Figure 2.**
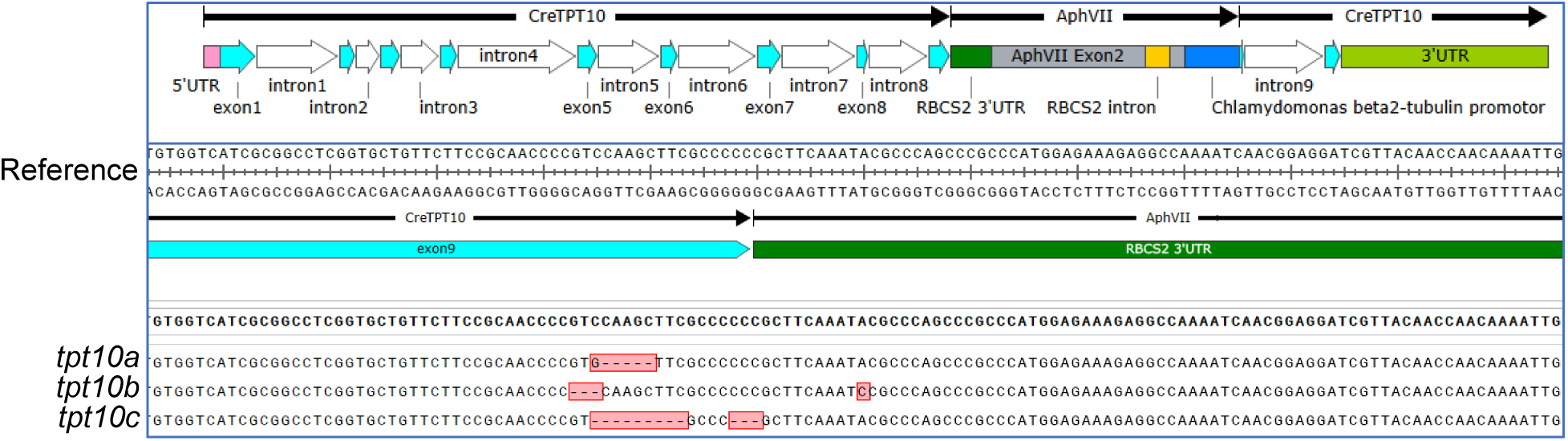
CRISPR/Cas9-mediated insertions in the *CreTPT10* gene. Sequencing of genomic DNA fragments across the site of insertion of the *AphVII* cassette in *tpt10a*, *tpt10b*, and *tpt10c*. The orientation and position of the *AphVII* cassette are presented above the alignment. The top sequence represents the expected cassette-genome junction reference sequence, constructed from the *CreTPT10* flanking genomic region and the *AphVII* cassette sequence at the insertion boundary. The mismatched nucleotides are marked in pink. Deletions are shown with dashes.

**Supplemental Figure 3.**
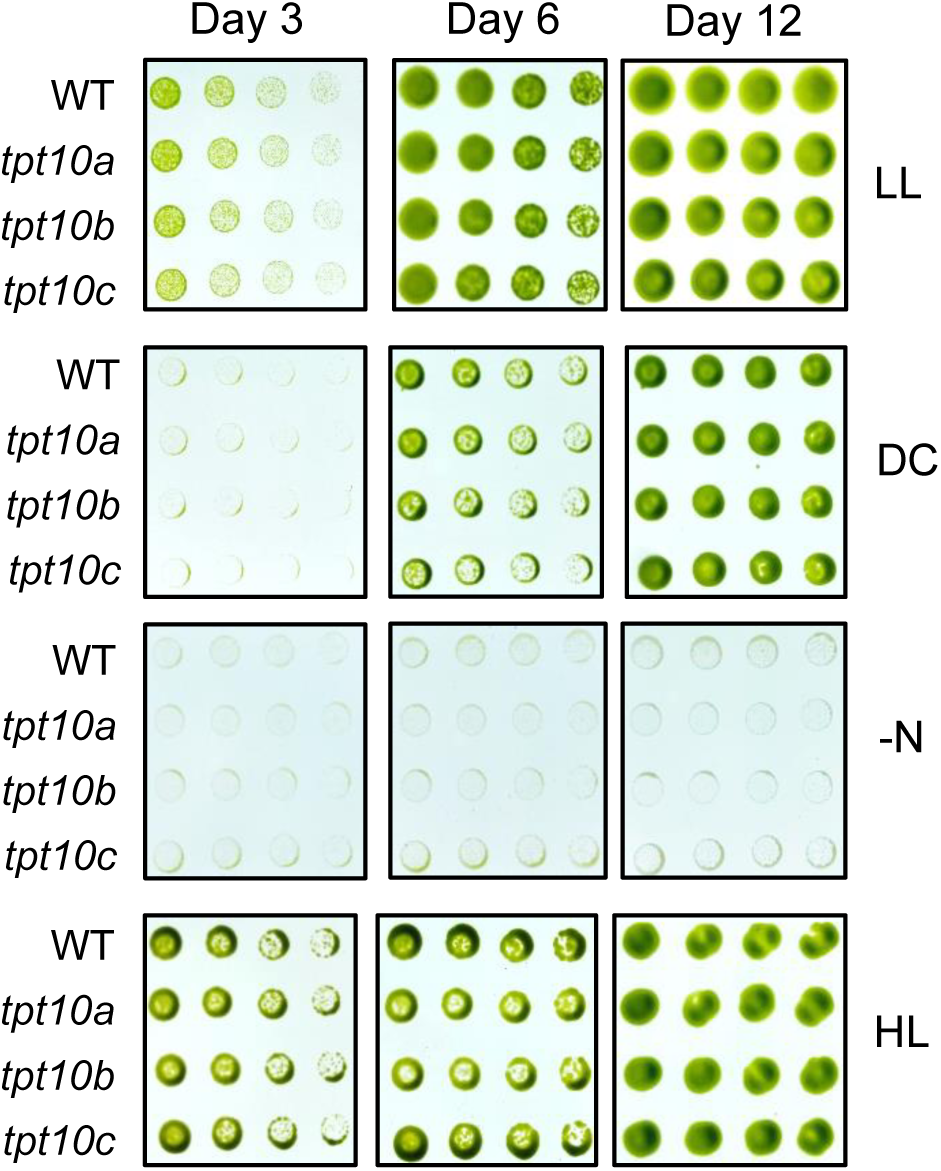
Growth analysis of WT and *tpt10* mutants on TAP agar plates. Growth of WT and *tpt10* mutants on TAP medium agar plates for 3, 6, and 12 days under LL, DC, −N, and HL. The dilution series was 4, 2, 1, and 0.5 μg mL^-1^ chlorophyll equivalent (left to right).

**Supplemental Figure 4.**
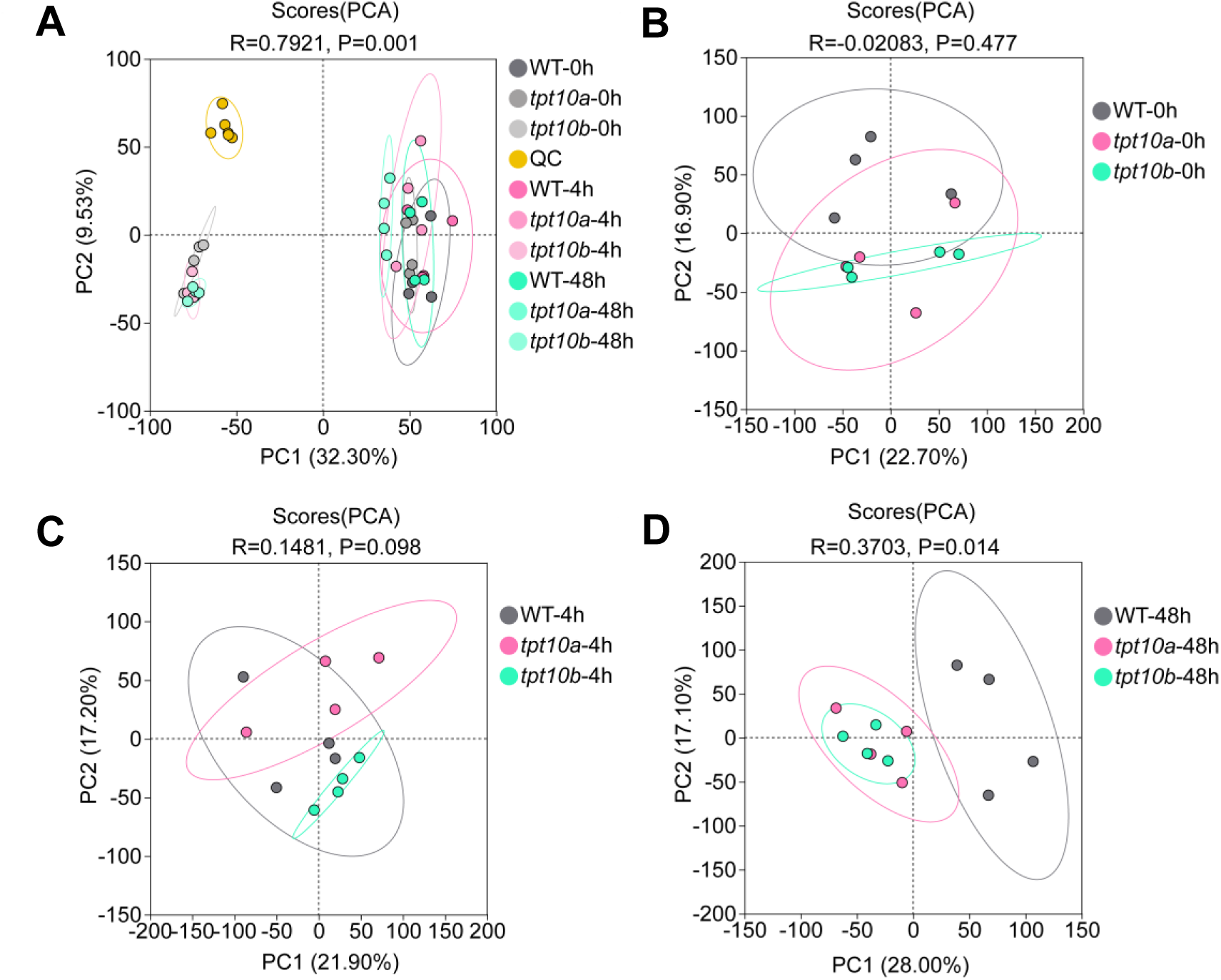
Principal component analysis (PCA) of metabolomic profiles in WT and *tpt10* mutants. **(A)** PCA scores for all samples across conditions (R = 0.7921, P = 0.001). **(B)** PCA scores under low light (R = −0.0208, P = 0.4). **(C)** PCA scores after 4 h of extended darkness (R = 0.1481, P = 0.098). **(D)** PCA scores after 48 h of extended darkness (R = 0.3703, P = 0.014).

**Supplemental Figure 5.**
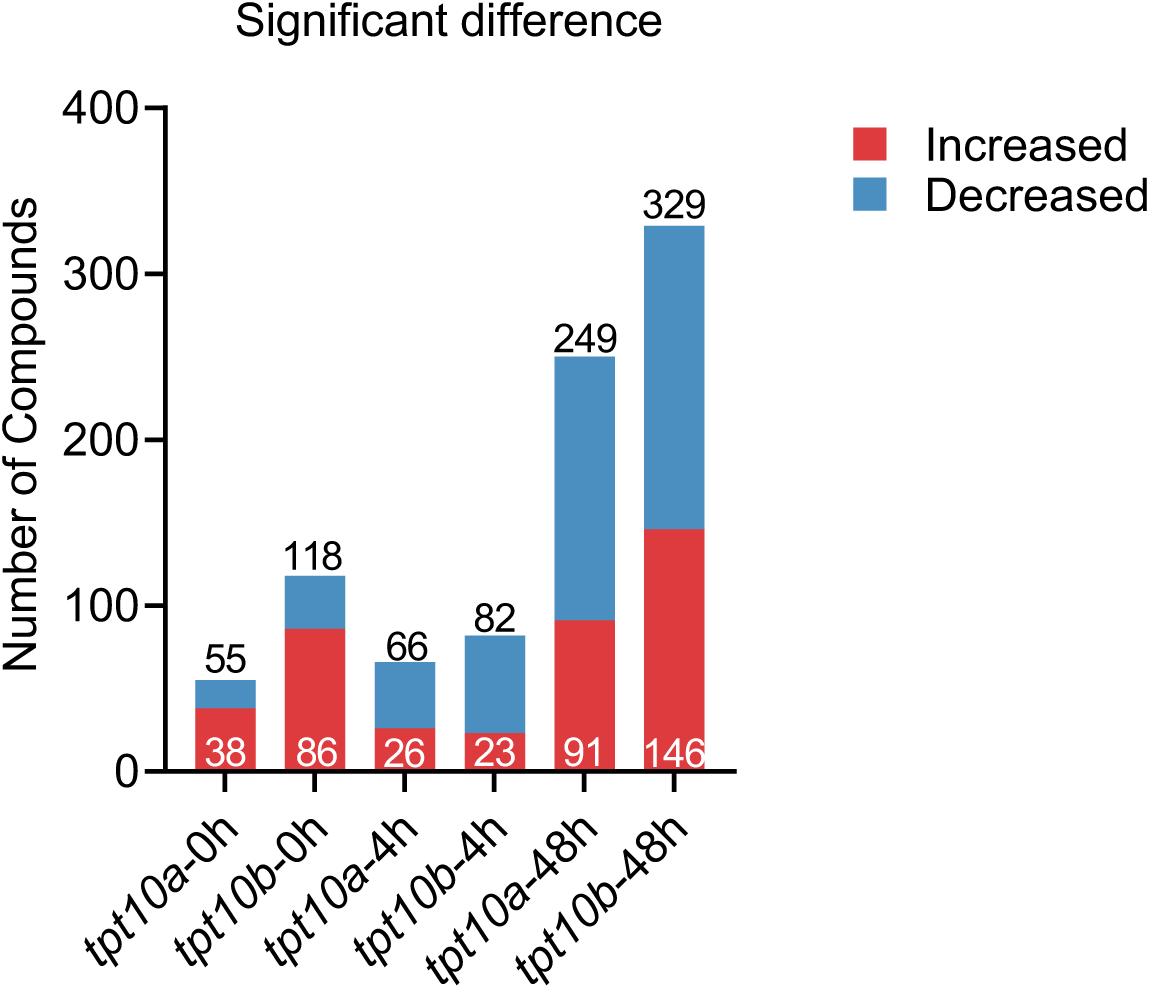
Numbers of compounds that change in abundance in *tpt10* mutants relative to WT cells during the transition from LL to extended darkness. Each stacked bar represents a mutant/WT comparison at the same time point (within-time-point differential analysis). Red and blue segments indicate the numbers of significantly increased and decreased compounds, respectively. Numbers within the red segments indicate the counts of increased compounds, and numbers above bars indicate the total numbers of significantly changed compounds.

**Supplemental Figure 6.**
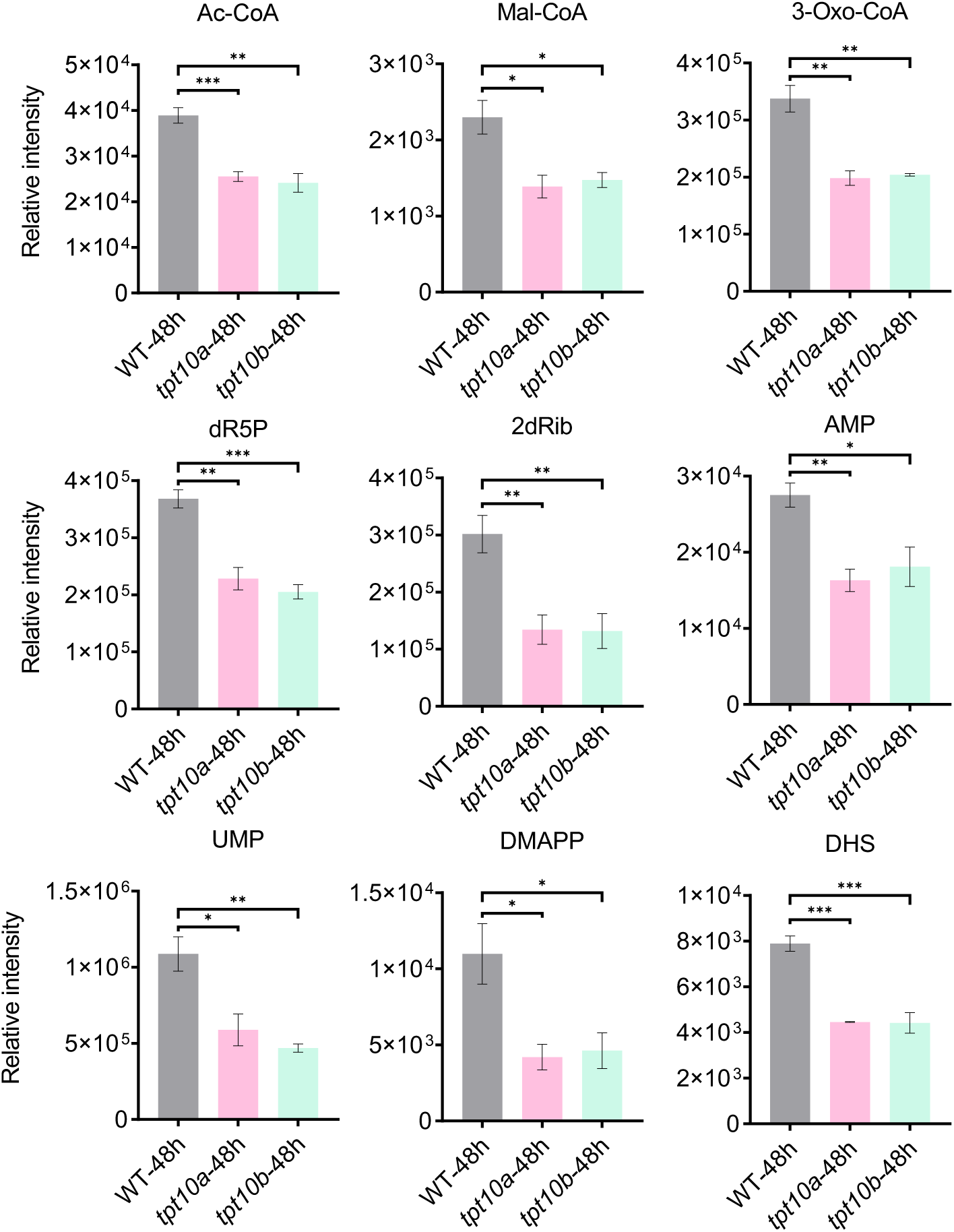
Depletion of plastid anabolism–associated metabolites in WT, *tpt10a*, and *tpt10b* after 48 h of extended darkness. Metabolite abundances were normalized to cell number and are shown as relative intensities. Bars represent means ± SEM (n = 4). Brackets indicate pairwise comparisons of each mutant versus WT using an unpaired two-tailed Student’s t-test (*P < 0.05, **P < 0.005, ***P < 0.001). Abbreviations: Ac-CoA, acetyl-CoA; Mal-CoA, malonyl-CoA; 3-Oxo-CoA, 3-oxooctanoyl-CoA; dR5P, deoxyribose 5-phosphate; 2dRib, 2-deoxy-D-ribose; AMP, adenosine monophosphate; UMP, uridine monophosphate; DMAPP, dimethylallyl diphosphate.

**Supplemental Figure 7.**
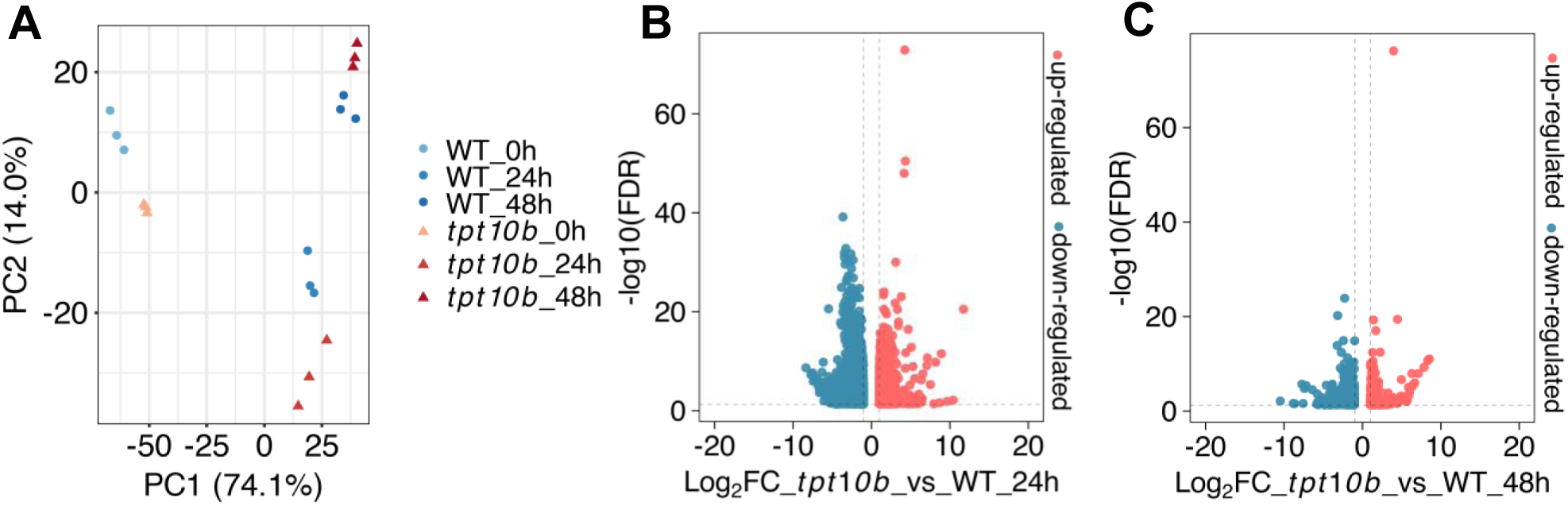
Transcriptomic divergence between WT and *tpt10b* during extended darkness. **(A)** PCA of transcriptomic profiles from WT and *tpt10b* strains at 0, 24, and 48 h of darkness. **(B and C)** Volcano plot of DEGs between *tpt10b* and WT at 24 h **(B)** and 48 h **(C)** of darkness. Red and blue dots indicate significantly up- and down-regulated genes, respectively (|log₂FC| > 1, P-adjust < 0.05).

**Supplemental Figure 8.**
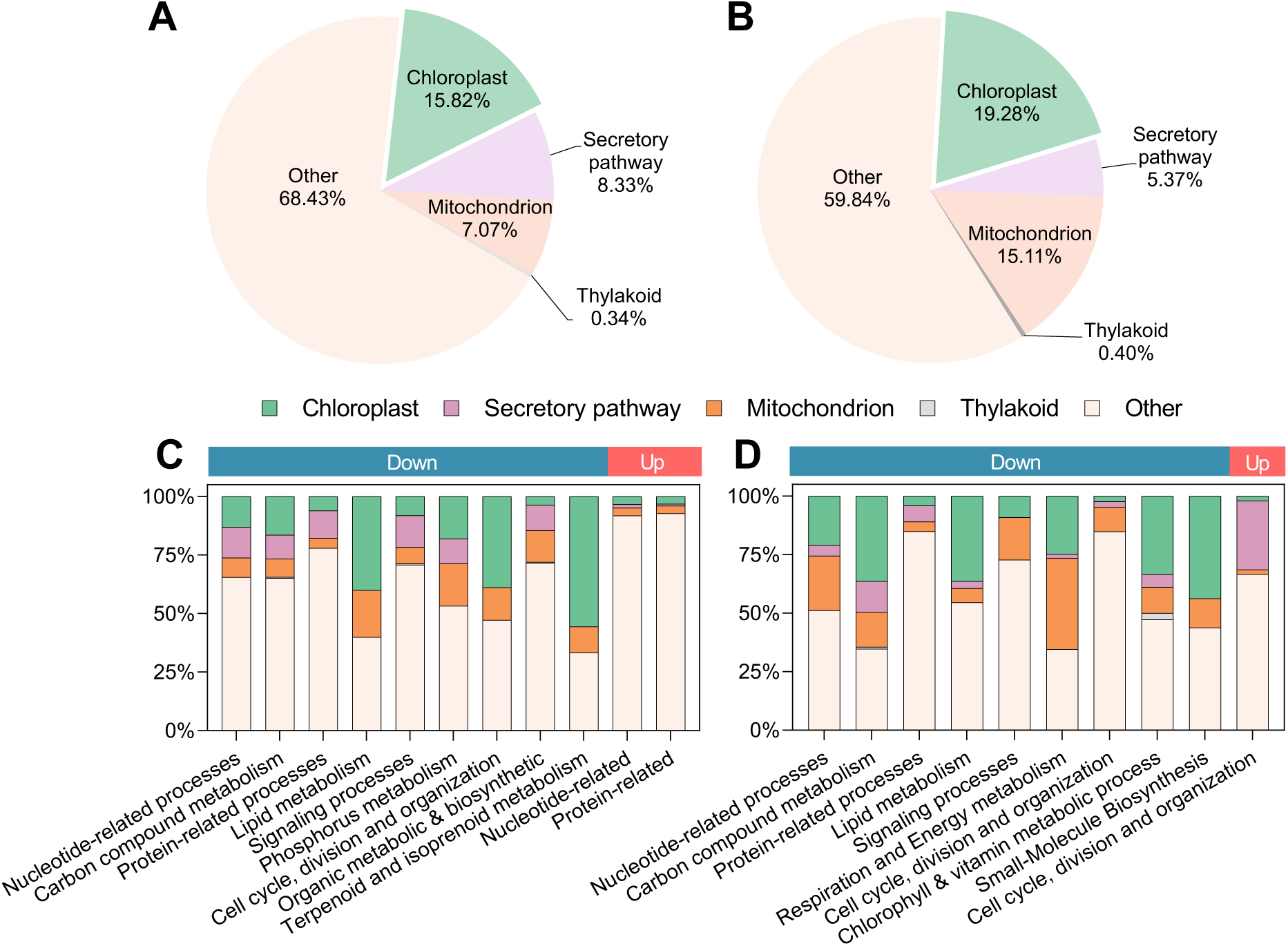
TargetP-predicted subcellular localization of proteins encoded by DEGs contributing to GO functional categories enriched under extended darkness. (A and. **B)** Pie charts of TargetP-predicted localization categories for DEGs contributing to GO functional categories enriched in the comparison of *tpt10* mutant vs WT after 24 h **(A)** and 48 h **(B)** of extended darkness. **(C and D)** Stacked bar charts of TargetP-predicted localization composition for DEGs within each enriched GO functional category after 24 h **(C)** and 48 h **(D)** of extended darkness. Percentages in pie charts indicate the fraction of DEGs assigned to each TargetP category (Chloroplast, Secretory pathway, Mitochondrion, Thylakoid, and Other). In stacked bar charts, each bar is normalized to 100% within a GO functional category and partitioned into the same TargetP categories. Other denotes proteins for which TargetP did not detect a canonical N-terminal targeting peptide signal, rather than a specific organelle localization. The color strip above the stacked bars indicates down-regulated genes (blue) and up-regulated genes (red).

**Supplemental Figure 9.**
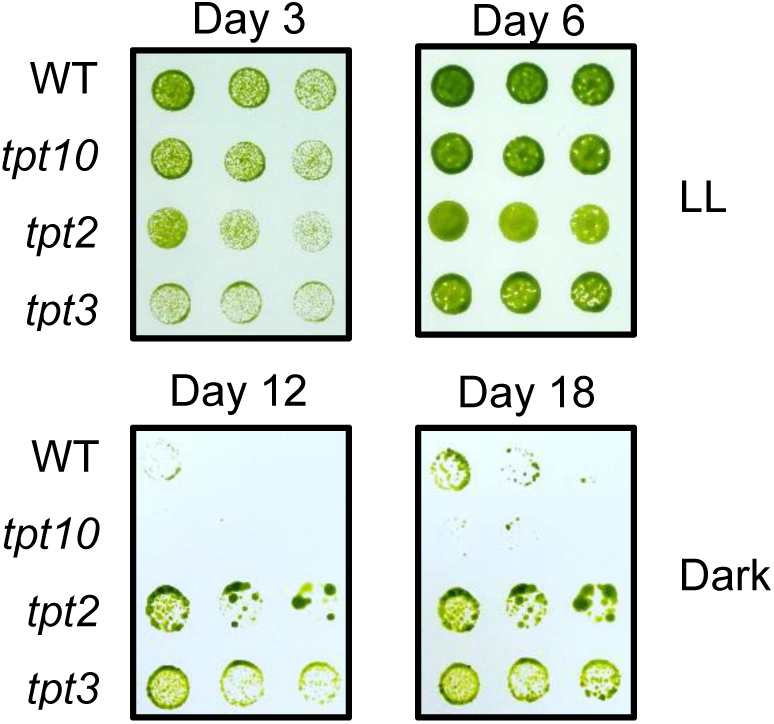
Growth analysis of WT and *pPTs* mutants on TAP agar plates. WT, *tpt10*, *tpt2*, and *tpt3* mutants were spotted onto TAP agar plates at serial chlorophyll concentrations of 4, 2, 1, and 0.5 μg mL^-1^ (left to right). Cultures were incubated under continuous low light (LL, 30 μmol photons m^-2^ s^-1^) for 3 and days, or under extended darkness (Dark, 0 μmol photons m^-2^ s^-1^) for 12 and 18 days.

**Supplemental Figure 10.**
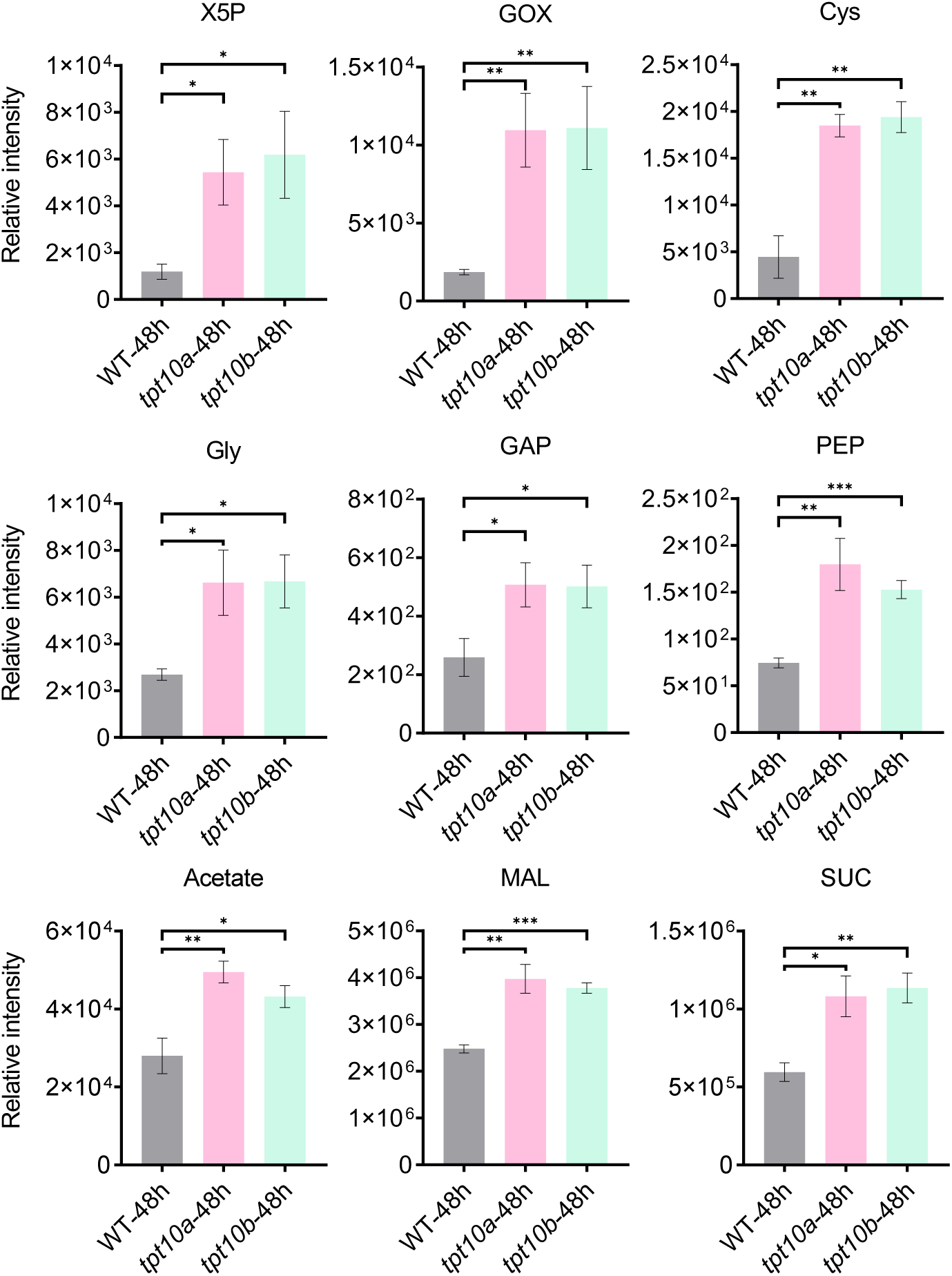
Accumulation of carbon intermediates and related amino acids in WT, *tpt10a*, and *tpt10b* after 48 h of extended darkness. Metabolite abundances were normalized to cell number and are shown as relative intensities. Bars represent means ± SEM (n = 4). Brackets indicate pairwise comparisons of each mutant versus WT using an unpaired two-tailed Student’s t-test (*P < 0.05, **P < 0.005, ***P < 0.001). Abbreviations: X5P, xylulose 5-phosphate; GOX, glyoxylate; Gly, glycine; Cys, cysteine; PEP, phosphoenolpyruvate; GAP, glyceraldehyde 3-phosphate; MAL, malate; SUC, succinate.

**Supplemental Figure 11.**
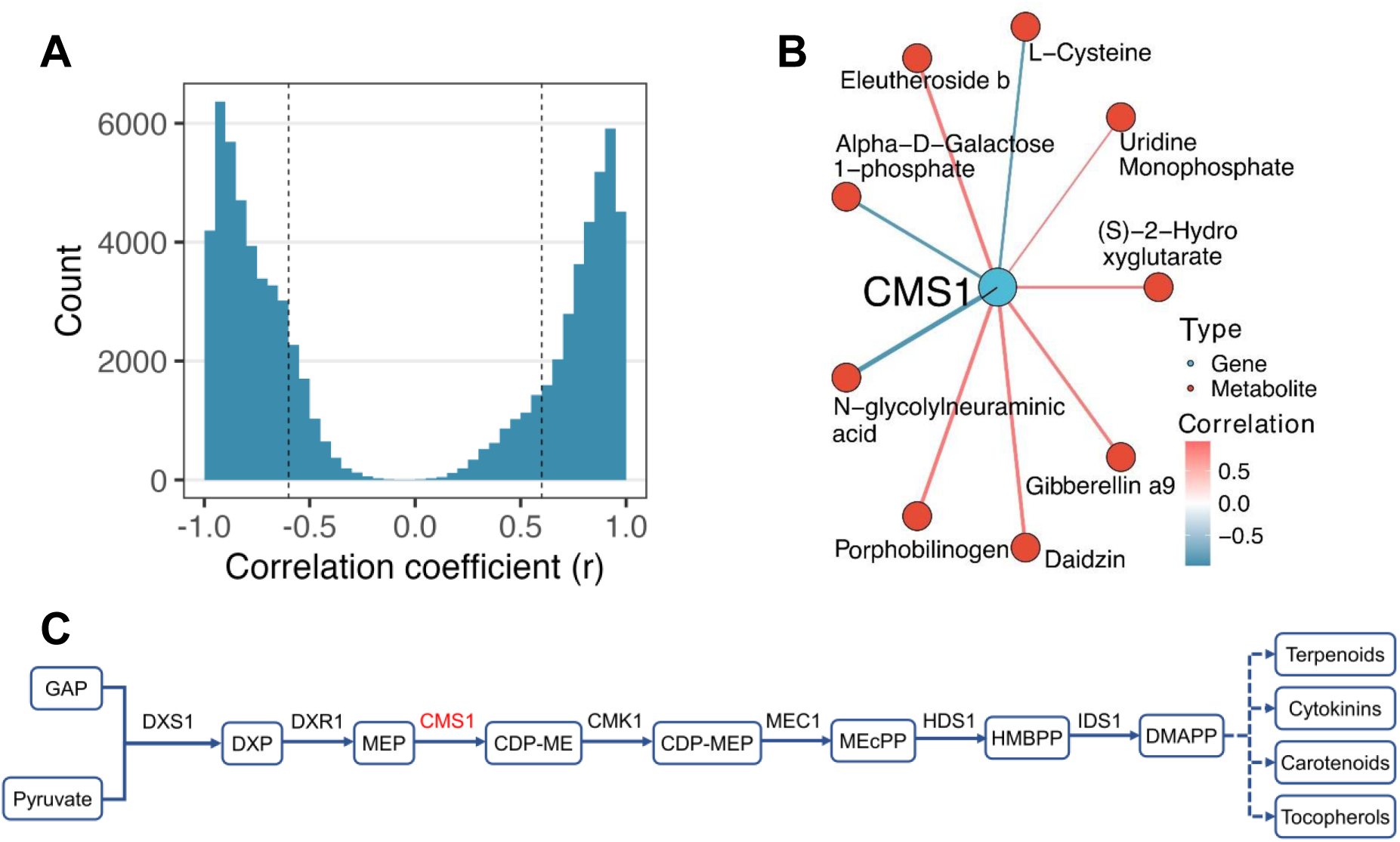
Integrated transcriptomic-metabolomic correlation analysis of 48 h extended darkness. **(A)** Distribution of metabolite-gene correlation coefficients calculated between significantly different metabolites and significantly different genes, with absolute values exceeding the threshold indicated by dashed lines (|r| cutoff), derived from integrated transcriptomic and metabolomic analyses at 48 h of darkness. **(B)** Representative correlation-based gene-metabolite network centered on CMS1, constructed using correlations between significantly different metabolites and significantly different genes. **(C)** Schematic overview of the MEP pathway showing CMS1 as a central enzyme within the pathway. Metabolite abbreviations: DXP, 1-deoxy-D-xylulose-5-phosphate; MEP, 2-C-methyl-D-erythritol 4-phosphate; CDP-ME, 4-(cytidine 5′-diphospho)-2-C-methyl-D-erythritol; CDP-MEP, 4-(cytidine 5′-diphospho)-2-C-methyl-D-erythritol 2-phosphate; MEcPP, 2-C-methyl-D-erythritol 2,4-cyclodiphosphate; HMBPP, (E)-4-hydroxy-3-methylbut-2-enyl diphosphate. Enzyme abbreviations: DXS, 1-deoxy-D-xylulose 5-phosphate synthase; DXR, 1-deoxy-D-xylulose-5-phosphate reductoisomerase; CMS1, 4-diphosphocytidyl-2C-methyl-D-erythritol synthase; CMK1, 4-diphosphocytidyl-2-C-methyl-D-erythritol kinase; MEC1, 2-C-methyl-D-erythritol 2,4-cyclodiphosphate synthase; HDS1, 1-hydroxy-2-methyl-2-(E)-butenyl 4-diphosphate synthase; IDS1, 4-hydroxy-3-methylbut-2-enyl diphosphate reductase.

**Supplemental Figure 12.**
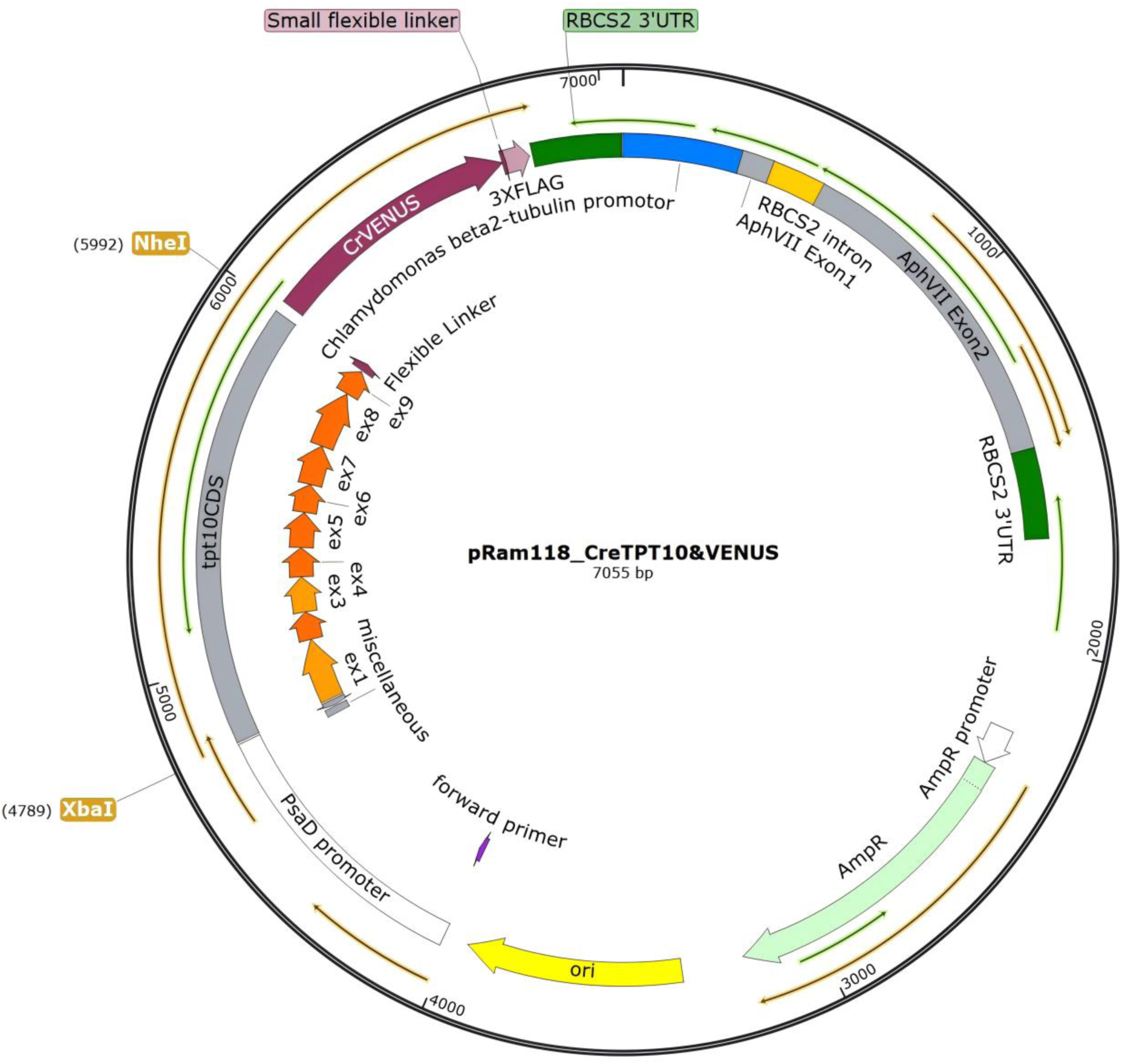
Map of the pRam118_*CreTPT10&VENUS* expression construct. Ram118_*CreTPT10&VENUS* (7,055 bp) contains the *PsaD* promoter driving full-length *CreTPT10* fused to *VENUS*, followed by the *RBCS2* 3’UTR. The *AphVII* hygromycin-resistance cassette is driven by the Chlamydomonas *β2-tubulin* promoter and followed by the *RBCS2* 3′UTR. The backbone vector pRAM118 was obtained from the Chlamydomonas Resource Center (sequence file provided in the CRC record) and is derived from pLM005 (GenBank KX077945.1). The CreTPT10 coding sequence corresponds to Cre08.g379350_4532 in the CC-4532 v6.1 reference genome (Phytozome/JGI).

**Supplemental Table 1.** List of differentially abundant metabolites (DAMs) annotated in Chlamydomonas after 48 h of extended darkness in *tpt10a* and *tpt10b* mutants relative to WT with fold changes and compound-class assignments, related to Figures 3 and 4.

**Supplemental Table 2.** List of DEGs with GO functional annotations and predicted subcellular localizations from TargetP and PB-Chlamy, related to Figures 5 and 6.

**Supplemental Table 3.**
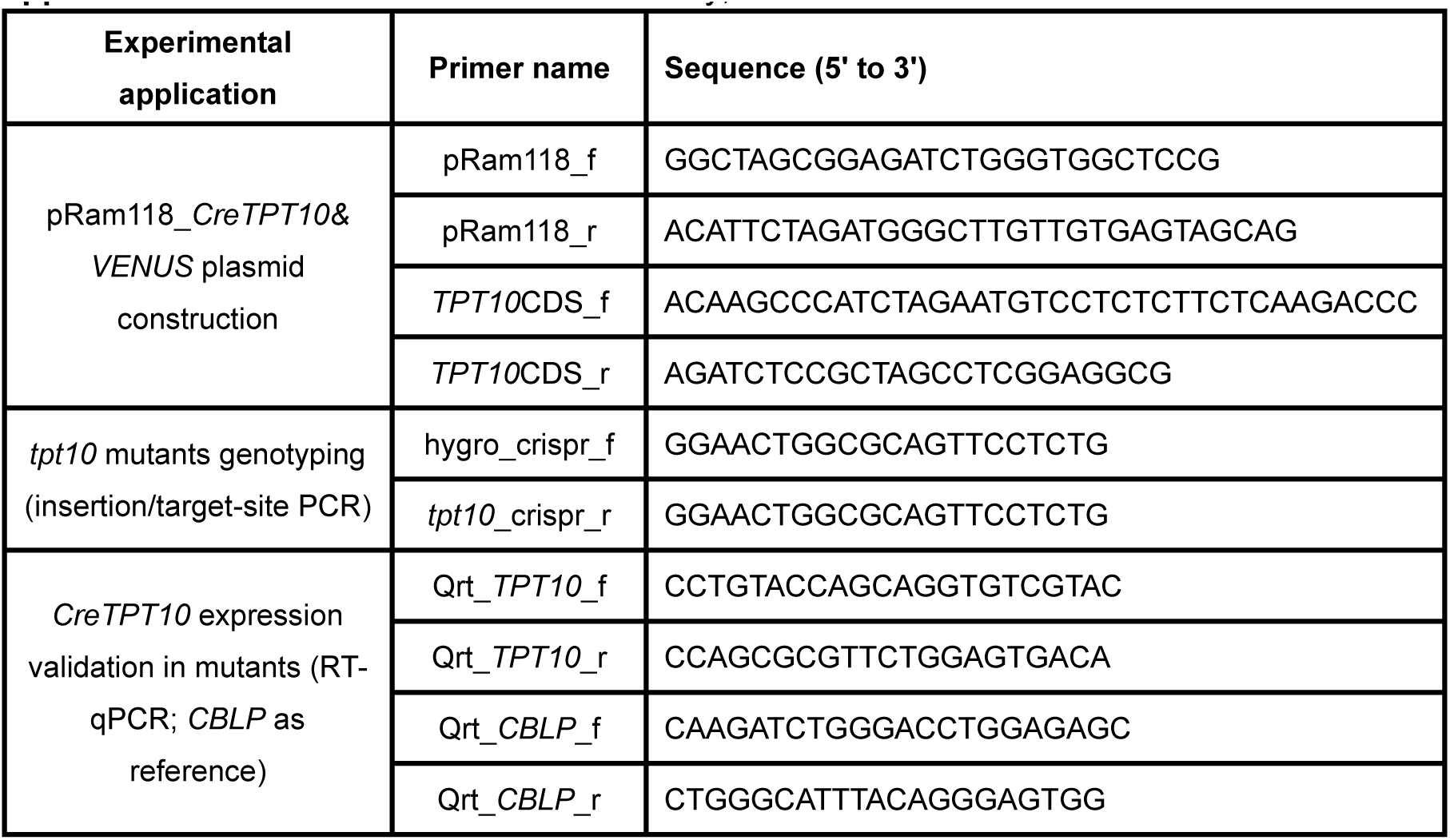
Primers used in this study, related to Methods.

**Supplemental Data Set 1.** The metabolome (cell–number–normalized peak intensity) for WT, *tpt10a*, and *tpt10b* populations across the transition from low light to extended darkness (0, 4, and 48 h) in four biological replicates.

**Supplemental Data Set 2.** Differentially abundant metabolites (DAMs) at 0 h (low light) and after 4 and 48 h of extended darkness in *tpt10a* vs WT and *tpt10b* vs WT comparisons, including DAMs shared between comparisons.

**Supplemental Data Set 3.** Significantly differentially expressed genes (DEGs) between WT and *tpt10b* at 24 and 48 h of extended darkness.

**Supplemental Data Set 4.** Gene Ontology (GO) enrichment of differentially expressed genes at 24 and 48 h of extended darkness.

## References

Almagro Armenteros, J. J., Salvatore, M., Emanuelsson, O., Winther, O., von Heijne, G., Elofsson, A., and Nielsen, H. (2019). Detecting sequence signals in targeting peptides using deep learning. Life Science Alliance 2:e201900429. 10.26508/lsa.201900429.

Ashburner, M., Ball, C. A., Blake, J. A., Botstein, D., Butler, H., Cherry, J. M., Davis, A. P., Dolinski, K., Dwight, S. S., Eppig, J. T., et al. (2000). Gene ontology: tool for the unification of biology. The gene ontology consortium. Nature Genetics 25:25–29. 10.1038/75556.

Blaby-Haas, C. E., Page, M. D., and Merchant, S. S. (2018). Using YFP as a Reporter of Gene Expression in the Green Alga *Chlamydomonas reinhardtii*. Methods in molecular biology (Clifton, N.J.) 1755:135–148. 10.1007/978-1-4939-7724-6_10.

Borowitzka, M. A. (2013). High-value products from microalgae—their development and commercialisation. Journal of Applied Phycology 25:743–756. 10.1007/s10811-013-9983-9.

Boyle, N. R., Page, M. D., Liu, B., Blaby, I. K., Casero, D., Kropat, J., Cokus, S. J., Hong-Hermesdorf, A., Shaw, J., Karpowicz, S. J., et al. (2012). Three Acyltransferases and Nitrogen-responsive Regulator Are Implicated in Nitrogen Starvation-induced Triacylglycerol Accumulation in *Chlamydomonas*. The Journal of Biological Chemistry 287:15811–15825. 10.1074/jbc.M111.334052.

Cheng, W.-H., Endo, A., Zhou, L., Penney, J., Chen, H.-C., Arroyo, A., Leon, P., Nambara, E., Asami, T., Seo, M., et al. (2002). A Unique Short-Chain Dehydrogenase/Reductase in Arabidopsis Glucose Signaling and Abscisic Acid Biosynthesis and Functions. The Plant Cell 14:2723–2743. 10.1105/tpc.006494.

Cho, M.-H., Lim, H., Shin, D. H., Jeon, J.-S., Bhoo, S. H., Park, Y.-I., and Hahn, T.-R. (2011). Role of the plastidic glucose translocator in the export of starch degradation products from the chloroplasts in *Arabidopsis thaliana*. New Phytologist 190:101–112. 10.1111/j.1469-8137.2010.03580.x.

Choi, B. Y., Shim, D., Kong, F., Auroy, P., Lee, Y., Li-Beisson, Y., Lee, Y., and Yamaoka, Y. (2022). The Chlamydomonas transcription factor MYB1 mediates lipid accumulation under nitrogen depletion. The New Phytologist 235:595–610. 10.1111/nph.18141.

Craig, R. J., Gallaher, S. D., Shu, S., Salomé, P. A., Jenkins, J. W., Blaby-Haas, C. E., Purvine, S. O., O’Donnell, S., Barry, K., Grimwood, J., et al. (2023). The Chlamydomonas genome project, version 6: reference assemblies for mating-type plus and minus strains reveal extensive structural mutation in the laboratory. Plant Cell 35:644–672. 10.1093/plcell/koac347.

DellaPenna, D., and Pogson, B. J. (2006). VITAMIN SYNTHESIS IN PLANTS: Tocopherols and Carotenoids. Annual Review of Plant Biology 57:711–738. 10.1146/annurev.arplant.56.032604.144301.

Doremus, H. D., and Jagendorf, A. T. (1985). Subcellular Localization of the Pathway of *de Novo* Pyrimidine Nucleotide Biosynthesis in Pea Leaves. Plant Physiology 79:856–861. 10.1104/pp.79.3.856.

Dupuis, S., Ojeda, V., Gallaher, S. D., Purvine, S. O., Glaesener, A. G., Ponce, R., Nicora, C. D., Bloodsworth, K., Lipton, M. S., Niyogi, K. K., et al. (2025). Too dim, too bright, and just right: systems analysis of the *Chlamydomonas* diurnal program under limiting and excess light. The Plant Cell 37:koaf086. 10.1093/plcell/koaf086.

Eicks, M., Maurino, V., Knappe, S., Flügge, U.-I., and Fischer, K. (2002). The Plastidic Pentose Phosphate Translocator Represents a Link between the Cytosolic and the Plastidic Pentose Phosphate Pathways in Plants. Plant Physiology 128:512–522. 10.1104/pp.010576.

Fan, J., Andre, C., and Xu, C. (2011). A chloroplast pathway for the de novo biosynthesis of triacylglycerol in *Chlamydomonas reinhardtii*. FEBS Letters 585:1985–1991. 10.1016/j.febslet.2011.05.018.

Findinier, J. (2023). Autolysin Production from *Chlamydomonas reinhardtii*. Bio-protocol 13. 10.21769/BioProtoc.4705.

Fischer, K., Kammerer, B., Gutensohn, M., Arbinger, B., Weber, A., Hausler, R. E., and Flugge, U. I. (1997). A new class of plastidic phosphate translocators: a putative link between primary and secondary metabolism by the phosphoenolpyruvate/phosphate antiporter. Plant Cell 9:453–462. 10.1105/tpc.9.3.453.

Flügge, U. I., Häusler, R. E., Ludewig, F., and Fischer, K. (2003). Functional genomics of phosphate antiport systems of plastids. Physiologia Plantarum 118:475–482. 10.1034/j.1399-3054.2003.00137.x.

Gibson, D. G., Young, L., Chuang, R.-Y., Venter, J. C., Hutchison, C. A., and Smith, H. O. (2009). Enzymatic assembly of DNA molecules up to several hundred kilobases. Nature Methods 6:343–345. 10.1038/nmeth.1318.

Gietz, R. D., and Schiestl, R. H. (2007). High-efficiency yeast transformation using the LiAc/SS carrier DNA/PEG method. Nature Protocols 2:31–34. 10.1038/nprot.2007.13.

Guijas, C., Montenegro-Burke, J. R., Domingo-Almenara, X., Palermo, A., Warth, B., Hermann, G., Koellensperger, G., Huan, T., Uritboonthai, W., Aisporna, A. E., et al. (2018). METLIN: A Technology Platform for Identifying Knowns and Unknowns. Analytical Chemistry 90:3156–3164. 10.1021/acs.analchem.7b04424.

Han, C., Shi, C., Liu, L., Han, J., Yang, Q., Wang, Y., Li, X., Fu, W., Gao, H., Huang, H., et al. (2024). Majorbio Cloud 2024: Update single-cell and multiomics workflows. iMeta 3:e217. 10.1002/imt2.217.

Häusler, R. E., Baur, B., Scharte, J., Teichmann, T., Eicks, M., Fischer, K. L., Flügge, U.-I., Schubert, S., Weber, A., and Fischer, K. (2000). Plastidic metabolite transporters and their physiological functions in the inducible crassulacean acid metabolism plant *mesembryanthemum crystallinum*. The Plant Journal 24:285–296. 10.1046/j.1365-313x.2000.00876.x.

Helliwell, C. A., Sullivan, J. A., Mould, R. M., Gray, J. C., Peacock, W. J., and Dennis, E. S. (2001). A plastid envelope location of *Arabidopsis* ent-kaurene oxidase links the plastid and endoplasmic reticulum steps of the gibberellin biosynthesis pathway. The Plant Journal: For Cell and Molecular Biology 28:201–208. 10.1046/j.1365-313x.2001.01150.x.

Herrmann, K. M., and Weaver, L. M. (1999). THE SHIKIMATE PATHWAY. Annual Review of Plant Physiology and Plant Molecular Biology 50:473–503. 10.1146/annurev.arplant.50.1.473.

Hilgers, E. J. A., Schöttler, M. A., Mettler-Altmann, T., Krueger, S., Dörmann, P., Eicks, M., Flügge, U.-I., and Häusler, R. E. (2018). The Combined Loss of Triose Phosphate and Xylulose 5-Phosphate/Phosphate Translocators Leads to Severe Growth Retardation and Impaired Photosynthesis in *Arabidopsis thaliana* tpt/xpt Double Mutants. Frontiers in Plant Science 9:1331. 10.3389/fpls.2018.01331.

Huang, W., Krishnan, A., Plett, A., Meagher, M., Linka, N., Wang, Y., Ren, B., Findinier, J., Redekop, P., Fakhimi, N., et al. (2023). *Chlamydomonas* mutants lacking chloroplast TRIOSE PHOSPHATE TRANSPORTER3 are metabolically compromised and light sensitive. Plant Cell 35:2592–2614. 10.1093/plcell/koad095.

Hung, W.-F., Chen, L.-J., Boldt, R., Sun, C.-W., and Li, H. (2004). Characterization of Arabidopsis Glutamine Phosphoribosyl Pyrophosphate Amidotransferase-Deficient Mutants. Plant Physiology 135:1314–1323. 10.1104/pp.104.040956.

Johnson, X., and Alric, J. (2013). Central Carbon Metabolism and Electron Transport in Chlamydomonas reinhardtii: Metabolic Constraints for Carbon Partitioning between Oil and Starch. Eukaryotic Cell 12:776–793. 10.1128/ec.00318-12.

Kamikawa, R., Moog, D., Zauner, S., Tanifuji, G., Ishida, K.-I., Miyashita, H., Mayama, S., Hashimoto, T., Maier, U. G., Archibald, J. M., et al. (2017). A Non-photosynthetic Diatom Reveals Early Steps of Reductive Evolution in Plastids. Molecular Biology and Evolution 34:2355–2366. 10.1093/molbev/msx172.

Kammerer, B., Fischer, K., Hilpert, B., Schubert, S., Gutensohn, M., Weber, A., and Flügge, U. I. (1998). Molecular Characterization of a Carbon Transporter in Plastids from Heterotrophic Tissues: The Glucose 6-Phosphate/Phosphate Antiporter. Plant Cell 10:105–117. 10.1105/tpc.10.1.105.

Kaye, Y., Huang, W., Clowez, S., Saroussi, S., Idoine, A., Sanz-Luque, E., and Grossman, A. R. (2019). The mitochondrial alternative oxidase from *Chlamydomonas reinhardtii* enables survival in high light. Journal of Biological Chemistry 294:1380–1395. 10.1074/jbc.RA118.004667.

Kim, D., Langmead, B., and Salzberg, S. L. (2015). HISAT: a fast spliced aligner with low memory requirements. Nature Methods 12:357–360. 10.1038/nmeth.3317.

Kumar, S., Stecher, G., Suleski, M., Sanderford, M., Sharma, S., and Tamura, K. (2024). MEGA12: Molecular Evolutionary Genetic Analysis Version 12 for Adaptive and Green Computing. Molecular Biology and Evolution 41:msae263. 10.1093/molbev/msae263.

Lanier, E. R., Andersen, T. B., and Hamberger, B. (2023). Plant terpene specialized metabolism: complex networks or simple linear pathways? The Plant journal: for Cell and Molecular Biology 114:1178–1201. 10.1111/tpj.16177.

Levasseur, W., Perré, P., and Pozzobon, V. (2020). A review of high value-added molecules production by microalgae in light of the classification. Biotechnology Advances 41:107545. 10.1016/j.biotechadv.2020.107545.

Li-Beisson, Y., Beisson, F., and Riekhof, W. (2015). Metabolism of acyl-lipids in *Chlamydomonas reinhardtii*. Plant Journal 82:504–522. 10.1111/tpj.12787.

Love, M. I., Huber, W., and Anders, S. (2014). Moderated estimation of fold change and dispersion for RNA-seq data with DESeq2. Genome Biology 15:550. 10.1186/s13059-014-0550-8.

Maeda, H., and Dudareva, N. (2012). The Shikimate Pathway and Aromatic Amino Acid Biosynthesis in Plants. Annual Review of Plant Biology 63:73–105. 10.1146/annurev-arplant-042811-105439.

Majeran, W., Zybailov, B., Ytterberg, A. J., Dunsmore, J., Sun, Q., and van Wijk, K. J. (2008). Consequences of C_4_ Differentiation for Chloroplast Membrane Proteomes in Maize Mesophyll and Bundle Sheath Cells. Molecular & Cellular Proteomics: MCP 7:1609–1638. 10.1074/mcp.M800016-MCP200.

Moog, D., Nozawa, A., Tozawa, Y., and Kamikawa, R. (2020). Substrate specificity of plastid phosphate transporters in a non-photosynthetic diatom and its implication in evolution of red alga-derived complex plastids. Scientific Reports 10:1167. 10.1038/s41598-020-58082-8.

Neuhaus, H. E., and Emes, M. J. (2000). NONPHOTOSYNTHETIC METABOLISM IN PLASTIDS. Annual Review of Plant Biology 51:111–140. 10.1146/annurev.arplant.51.1.111.

Niewiadomski, P., Knappe, S., Geimer, S., Fischer, K., Schulz, B., Unte, U. S., Rosso, M. G., Ache, P., Flügge, U.-I., and Schneider, A. (2005). The Arabidopsis Plastidic Glucose 6-Phosphate/Phosphate Embryo Translocator GPT, Is Essential for Pollen Maturation and Sac Development. The Plant Cell 17:760–775. 10.1105/tpc.104.029124.

Niittylä, T., Messerli, G., Trevisan, M., Chen, J., Smith, A. M., and Zeeman, S. C. (2004). A Previously Unknown Maltose Transporter Essential for Starch Degradation in Leaves. Science 303:87–89. 10.1126/science.1091811.

Ohlrogge, J., and Browse, J. (1995). Lipid biosynthesis. The Plant Cell 7:957–970. 10.1105/tpc.7.7.957.

Porra, R. J., Thompson, W. A., and Kriedemann, P. E. (1989). Determination of accurate extinction coefficients and simultaneous equations for assaying chlorophylls a and b extracted with four different solvents: verification of the concentration of chlorophyll standards by atomic absorption spectroscopy. Biochimica et Biophysica Acta (BBA) - Bioenergetics 975:384–394. 10.1016/S0005-2728(89)80347-0.

Pott, D. M., Osorio, S., and Vallarino, J. G. (2019). From Central to Specialized Metabolism: An Overview of Some Secondary Compounds Derived From the Primary Metabolism for Their Role in Conferring Nutritional and Organoleptic Characteristics to Fruit. Frontiers in Plant Science 10. 10.3389/fpls.2019.00835.

Rolletschek, H., Nguyen, T. H., Häusler, R. E., Rutten, T., Göbel, C., Feussner, I., Radchuk, R., Tewes, A., Claus, B., Klukas, C., et al. (2007). Antisense inhibition of the plastidial glucose-6-phosphate/phosphate translocator in *vicia* seeds shifts cellular differentiation and promotes protein storage. Plant Journal 51:468–484. 10.1111/j.1365-313X.2007.03155.x.

Romer, J., Gutbrod, K., Schuppener, A., Melzer, M., Müller-Schüssele, S. J., Meyer, A. J., and Dörmann, P. (2024). Tocopherol and phylloquinone biosynthesis in chloroplasts requires the phytol kinase VITAMIN E PATHWAY GENE5 (VTE5) and the farnesol kinase (FOLK). The Plant Cell 36:1141–1158. 10.1093/plcell/koad316.

Ruiz-Sola, M. Á., and Rodríguez-Concepción, M. (2012). Carotenoid Biosynthesis in Arabidopsis: A Colorful Pathway. The Arabidopsis Book 10:e0158. 10.1199/tab.0158.

Sakakibara, H. (2025). Five unaddressed questions about cytokinin biosynthesis. Journal of Experimental Botany 76:1941–1949. 10.1093/jxb/erae348.

Schmid, J., and Amrhein, N. (1995). Molecular organization of the shikimate pathway in higher plants. Phytochemistry 39:737–749. 10.1016/0031-9422(94)00962-S.

Shumskaya, M., Bradbury, L. M. T., Monaco, R. R., and Wurtzel, E. T. (2012). Plastid Localization of the Key Carotenoid Enzyme Phytoene Synthase Is Altered by Isozyme, Allelic Variation, and Activity. The Plant Cell 24:3725–3741. 10.1105/tpc.112.104174.

Silver, N., Best, S., Jiang, J., and Thein, S. L. (2006). Selection of housekeeping genes for gene expression studies in human reticulocytes using real-time PCR. BMC Molecular Biology 7:33. 10.1186/1471-2199-7-33.

Staehr, P., Löttgert, T., Christmann, A., Krueger, S., Rosar, C., Rolčík, J., Novák, O., Strnad, M., Bell, K., Weber, A. P. M., et al. (2014). Reticulate leaves and stunted roots are independent phenotypes pointing at opposite roles of the phosphoenolpyruvate/phosphate translocator defective in *cue1* in the plastids of both organs. Frontiers in Plant Science 5. 10.3389/fpls.2014.00126.

Streatfield, S. J., Weber, A., Kinsman, E. A., Häusler, R. E., Li, J. M., Post-Beittenmiller, D., Kaiser, W. M., Pyke, K. A., Flügge, U. I., and Chory, J. (1999). The Phosphoenolpyruvate/Phosphate Translocator Is Required for Phenolic Metabolism, Palisade Cell Development, and Plastid-Dependent Nuclear Gene Expression. Plant Cell 11:1609–1621. 10.1105/tpc.11.9.1609.

Sussulini, A. ed. (2017). Metabolomics: From Fundamentals to Clinical Applications. Cham: Springer International Publishing. 10.1007/978-3-319-47656-8

Takei, K., Yamaya, T., and Sakakibara, H. (2004). *Arabidopsis* CYP735A1 and CYP735A2 Encode Cytokinin Hydroxylases That Catalyze the Biosynthesis of trans-Zeatin. Journal of Biological Chemistry 279:41866–41872. 10.1074/jbc.M406337200.

Tanaka, R., and Tanaka, A. (2007). Tetrapyrrole Biosynthesis in Higher Plants. Annual Review of Plant Biology 58:321–346. 10.1146/annurev.arplant.57.032905.105448.

Tang, S., Peng, F., Tang, Q., Liu, Y., Xia, H., Yao, X., Lu, S., and Guo, L. (2022). BnaPPT1 is essential for chloroplast development and seed oil accumulation in Brassica napus. Journal of Advanced Research 42:29–40. 10.1016/j.jare.2022.07.008.

Vranová, E., Coman, D., and Gruissem, W. (2013). Network Analysis of the MVA and MEP Pathways for Isoprenoid Synthesis. Annual Review of Plant Biology 64:665–700. 10.1146/annurev-arplant-050312-120116.

Wang, L., Patena, W., Baalen, K. A. V., Xie, Y., Singer, E. R., Gavrilenko, S., Warren-Williams, M., Han, L., Harrigan, H. R., Hartz, L. D., et al. (2023). A chloroplast protein atlas reveals punctate structures and spatial organization of biosynthetic pathways. Cell 186:3499–3518.e14. 10.1016/j.cell.2023.06.008.

Warakanont, J., Tsai, C.-H., Michel, E. J. S., Murphy III, G. R., Hsueh, P. Y., Roston, R. L., Sears, B. B., and Benning, C. (2015). Chloroplast lipid transfer processes in *Chlamydomonas reinhardtii* involving a TRIGALACTOSYLDIACYLGLYCEROL 2 (TGD2) orthologue. The Plant Journal 84:1005–1020. 10.1111/tpj.13060.

Weber, A. P. M., Schneidereit, J., and Voll, L. M. (2004). Using mutants to probe the *in vivo* function of plastid envelope membrane metabolite transporters. Journal of Experimental Botany 55:1231–1244. 10.1093/jxb/erh091.

Weber, A. P. M., Schwacke, R., and Flügge, U.-I. (2005). SOLUTE TRANSPORTERS OF THE PLASTID ENVELOPEMEMBRANE. Annual Review of Plant Biology 56:133–164. 10.1146/annurev.arplant.56.032604.144228.

Wishart, D. S., Guo, A., Oler, E., Wang, F., Anjum, A., Peters, H., Dizon, R., Sayeeda, Z., Tian, S., Lee, B. L., et al. (2022). HMDB 5.0: the Human Metabolome Database for 2022. Nucleic Acids Research 50:D622–D631. 10.1093/nar/gkab1062.

Witte, C.-P., and Herde, M. (2020). Nucleotide Metabolism in Plants. Plant Physiology 182:63–78. 10.1104/pp.19.00955.

Witz, S., Jung, B., Fürst, S., and Möhlmann, T. (2012). De Novo Pyrimidine Nucleotide Synthesis Mainly Occurs outside of Plastids, but a Previously Undiscovered Nucleobase Importer Provides Substrates for the Essential Salvage Pathway in *Arabidopsis*. The Plant Cell 24:1549–1559. 10.1105/tpc.112.096743.

Yang, Z. (1994). Maximum likelihood phylogenetic estimation from DNA sequences with variable rates oversites: Approximate methods. Journal of Molecular Evolution 39:306–314. 10.1007/BF00160154.

Zhao, M.-L., Cai, W.-S., Zheng, S.-Q., Zhao, J.-L., Zhang, J.-L., Huang, Y., Hu, Z.-L., and Jia, B. (2022). Metabolic Engineering of the Isopentenol Utilization Pathway Enhanced the Production of Terpenoids in *Chlamydomonas reinhardtii*. Marine Drugs 20:577. 10.3390/md20090577.

Zheng, Y., Deng, X., Qu, A., Zhang, M., Tao, Y., Yang, L., Liu, Y., Xu, J., and Zhang, S. (2018). Regulation of pollen lipid body biogenesis by MAP kinases and downstream WRKY transcription factors in *Arabidopsis*. PLOS Genetics 14:e1007880. 10.1371/journal.pgen.1007880.

Zrenner, R., Stitt, M., Sonnewald, U., and Boldt, R. (2006). PYRIMIDINE AND PURINE BIOSYNTHESIS AND DEGRADATION IN PLANTS. Annual Review of Plant Biology 57:805–836. 10.1146/annurev.arplant.57.032905.105421.

